# A distinct class of conjugative megaplasmids includes potential vehicles for prophage dissemination

**DOI:** 10.64898/2026.02.21.707213

**Authors:** Ling Yuan, Yiting Qin, Jacob West-Roberts, Karthik Anantharaman, Haoyu Wang, Yuanqiang Zou, Yi Duan, Antonio Pedro Camargo, Eugene V. Koonin, LinXing Chen

**Affiliations:** State Key Laboratory of Advanced Environmental Technology, the Department of Environmental Science and Engineering, University of Science and Technology of China, Hefei, China; Innovative Genomics Institute, University of California, Berkeley, Berkeley, CA 94720, USA; Department of Bacteriology, University of Wisconsin-Madison, Madison, WI, USA; State Key Laboratory of Genome and Multi-omics Technologies, BGI Research, Shenzhen 518083, China; College of Life Sciences, University of Chinese Academy of Sciences, Beijing 100049, China; State Key Laboratory of Immune Response and Immunotherapy, Department of Infectious Diseases, The First Affiliated Hospital of USTC, Center for Advanced Interdisciplinary Science and Biomedicine of IHM, Division of Life Sciences and Medicine, University of Science and Technology of China, Hefei, Anhui 230027, China; Key Laboratory of Anhui Province for Emerging and Reemerging Infectious Diseases, Hefei, Anhui 230027, China; Department of Biochemistry, Institute of Chemistry, University of São Paulo, São Paulo, SP 05508-060, Brazil; Computational Biology Branch, Division of Intramural Research, National Library of Medicine, National Institutes of Health, Bethesda, MD 20894, USA

**Keywords:** Prophage, Plasmid, Lysogeny, Human gut, Metagenomics, Conjugative transfer

## Abstract

Closely related prophages are frequently found in phylogenetically distant bacteria in the human gut, despite limited evidence of productive phage infections across broad host ranges. Thus, it remains unclear how the wide distribution of prophages could emerge. Here, we identify a potential mechanism of prophage dissemination. We describe two deeply diverged groups of conjugative megaplasmids (>300 kilobases) in the human gut microbiome, which we term Hodors. Hodors encode conserved replication, partitioning, and type IV secretion systems, together with a complex surface-associated gene module. A subset of Hodors harbor complete, intact prophage genomes, and closely related prophages are detected across phylogenetically distant Bacillota lineages, including both Bacilli and Clostridia. Further analysis indicates that Hodor-associated prophages can exist as extracellular particles and demonstrate their transcriptional activity. Our findings support a model in which conjugative megaplasmids act as composite mobile platforms that disseminate prophage genomes across bacterial lineages, providing a mechanistic explanation for the widespread occurrence of closely related prophages in phylogenetically distant gut bacteria and effectively decoupling lysogenic host range from infective host range.

## Introduction

Viruses are the most abundant biological entities on Earth, with virus particles outnumbering cells several-fold in many habitats ^1–3^. Phages are integral components of microbial ecosystems and play central roles in shaping bacterial population dynamics, genome evolution, and horizontal gene transfer ^4^. Experimental infection assays have consistently shown that most phages have narrow host ranges, typically restricted to the bacterial strain or species level ^5^, and on some occasions, to different species from different genera in the same family ^6–8^, whereas infection across bacterial families is extremely rare. Long-term persistence of phages is commonly achieved through lysogeny within a limited set of related hosts ^9,10^. Numerous studies of the human gut microbiome have reported that the majority of prophages rarely enter productive lytic cycles and instead remain stably integrated within host genomes ^11,12^. However, culture-independent analyses have revealed an apparent contradiction, with several recent studies reporting that closely related prophages are frequently found integrated into genomes of phylogenetically distant bacteria (cross-family) in the human gut ^13–15^. This pattern challenges the classical views of phage host specificity and raises a fundamental question: how can temperate phage genomes become broadly distributed across divergent hosts despite limited evidence of productive infection in the human gut?

Here, we report a previously unrecognized mechanism that can provide an explanation for this phenomenon. We identify two deeply diverged groups of conjugative megaplasmids exceeding 300 kbp in size in the human gut microbiome, which we term Hodors. A subset of these megaplasmids carries complete, intact prophage genomes, some of which are integrated in genomes of distantly related Bacillota lineages, including both Bacilli and Clostridia, while exhibiting only limited genomic divergences. These observations suggest that temperate phage genomes can be disseminated across the bacterial diversity via plasmid-mediated transfer and subsequently integrated into the host chromosomes, without requiring the capability to infect an unusually broad range of hosts. Thus, our findings establish conjugative megaplasmids as vehicles for temperate phage dispersal and provide a plausible mechanistic framework for understanding how broadly distributed prophages can arise in the human gut and other microbiomes.

## Results

### Hodors are megaplasmids inhabiting the human gut, with some harbouring complete, intact prophage genomes

In our analysis of the huge phage genome collection (HPGC) ^16^, we noticed that within two groups of genomes, only two genomes in each group encoded a large terminase subunit (TerL), a conserved, hallmark gene present in all tailed phages, that was identified using sequence and structural searches, although paired-end reads mapping indicated that some genomes lacking TerL genes were circular (Supplementary Table 1). These two groups of genomes were defined at ≥90% nucleotide identity and across ≥80% genome length, and consisted of 9 (G1) and 10 (G2) genomes, respectively. These genomes were 211-389 kilobase pairs (kbp) in size and were all reconstructed from human gut metagenomic samples ^17–19^. Gene prediction and annotation confirmed that they encoded canonical plasmid functions, including a replication module with an initiation protein (Rep_3) and partitioning proteins such as ParA, ParB, and ParM/StbA (Supplementary Table 2). Thus, we concluded that these genomes represent megaplasmids. Notably, all of 19 genomes encoded multiple sortases, which are transpeptidases that covalently anchor surface proteins to the cell envelope in Gram-positive bacteria ^20^. We designate this set of megaplasmids “Hodors” hereafter as an operational label reflecting their shared genomic features and unusually large size. The name is inspired by the fictional character Hodor from A Song of Ice and Fire, known for “holding the door,” and is used here as a mnemonic reference to their large and cohesive genomic structure.

Manual genome curation showed that one of the TerL genes was associated with a Hodor due to chimeric assembly (Supplementary Figure 1), but the other three genomes (Hodor01, 02, and 10) each contained a complete prophage genome (Supplementary Figure 2). Using Pebblescout search ^21^ and *do novo* assembly, we sought to curate additional Hodor genomes (Methods). As a result, we obtained a total of 38 high-quality Hodor genomes (25 of these were complete), which had an average length of 356.8 kbp (Supplementary Table 1). Among them, 11 fell within G1, and 27 belonged to G2, with G1 genomes having a slightly higher GC content than G2 ones (47.1% vs 44.6%).

We compared the protein sequences encoded by the Hodors to the UniProtKB database and found that >35% of these were most similar to homologs from Clostridia and Bacilli, the two large, well-characterized classes of the phylum Bacillota (previously, Firmicutes) (Figure 1a), for example, the class B sortases (Supplementary Figure 3, Supplementary Table 3), suggesting these bacteria could be the hosts of Hodors. Although we removed the prophage genomes from the Hodors before conducting the database search, phage-related proteins accounted for 3.3% of all hits, on average (2.5-4.4%). Among the phage-related proteins, TnpB nucleases (actually, encoded by insertion sequence, that is, transposable elements integrated in phage genomes) and Phage_lysozyme2 domains are frequently identified (Supplementary Table 4).

**Figure 1.**
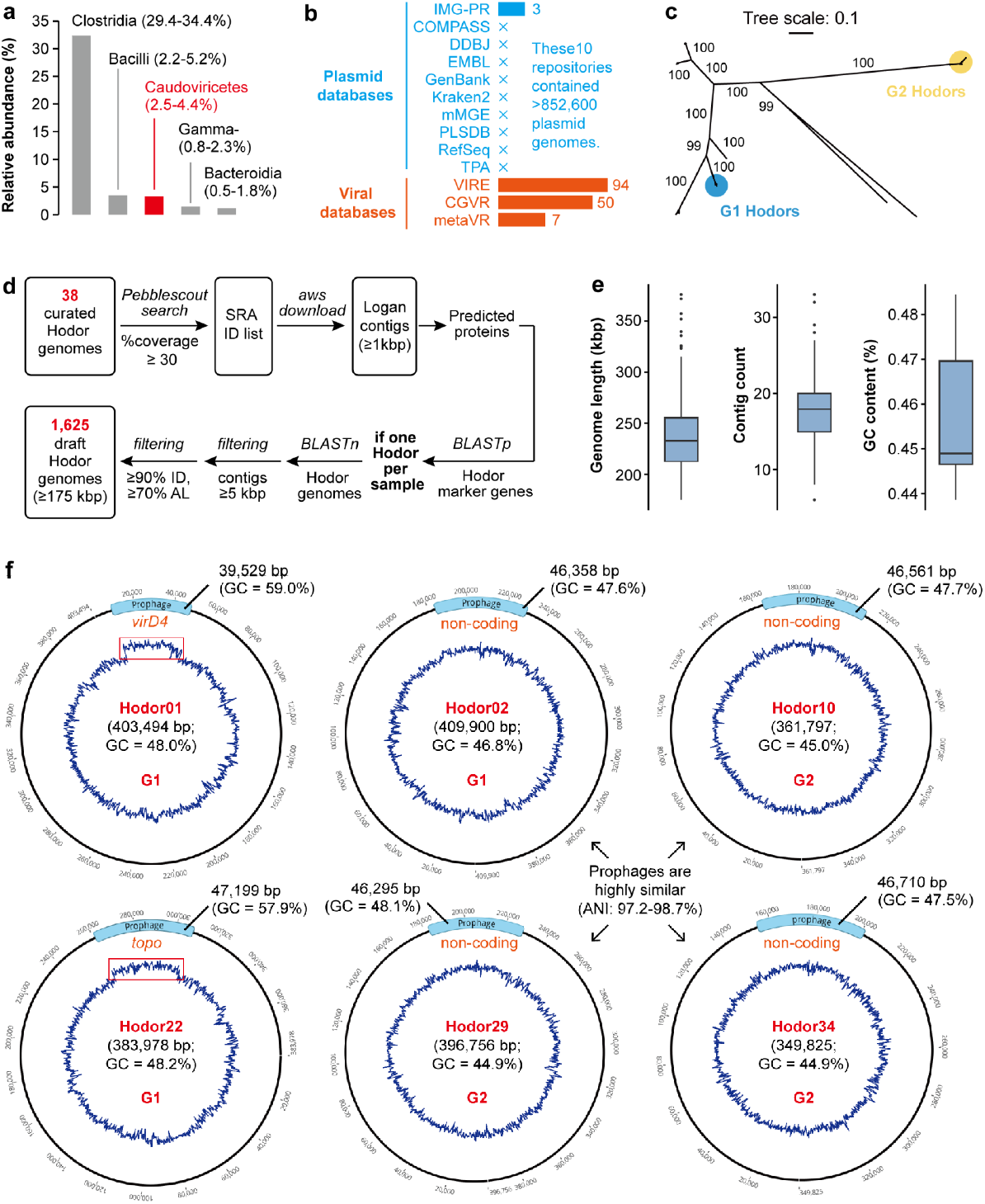
Identification of circular megaplasmid genomes with embedded complete prophage genomes. (a) Taxonomic affiliations of the closest UniProtKB homologs of Hodor-encoded genes. The five most frequent classes are displayed. (b) Detection of Hodors and related elements in viral and plasmid databases. Bars indicate log10-scaled counts. Proteins predicted from plasmid genomes were searched against four Hodor replication and partitioning proteins (Rep_3, ParA, ParB, and ParM/StbA) using BLASTp (≥50% identity over ≥70% length). Plasmids matching at least one marker were considered Hodor relatives. (c) Phylogeny of Hodors and relatives based on concatenated Rep_3, ParA, and ParB sequences, with bootstrap values shown. (d) Pipeline for recovering Hodor draft genomes from Logan assemblies. Curated Hodor genomes were queried against Pebblescout to identify non-redundant metagenomic samples, from which contigs were retrieved and compared to the curated Hodor genomes. Draft genomes were defined as assemblies >175 kbp, corresponding to ≥50% completeness relative to the average size (351 kbp) of 25 curated complete genomes (prophages excluded). (e) Summary statistics of Logan-derived draft Hodor genomes, including genome length, contig number, and GC content. (f) The 6 complete megaplasmid genomes with embedded complete prophage genomes. The divergent GC profiles of the prophage regions in Hodor01 and Hodor22 are highlighted with red boxes. The prophage genomes in Hodor02, Hodor10, Hodor29, and Hodor34 have an average nucleotide identity (ANI) of 97.2-98.7%. The locations and the information of the prophage regions are shown. The genes in which the prophages were inserted are shown in orange. *topo*, DNA topoisomerase.

Among the 38 curated Hodor genomes, 6 (three from each group) harboured complete, intact prophages (Figure 1f). Notably, although Hodor01 and Hodor02 were both from G1, they contained different prophages inserted at different positions in their genomes (39,529 vs 46,358 bp in length), with TerL being 12.5% identity. Conversely, despite Hodor02 from G1 being distantly related to Hodor10, Hodor29, and Hodor34 from G2, all these Hodors shared closely similar prophage genomes with 97.2-98.7% average nucleotide identity (ANI) (Figure 1f, Supplementary Figure 5), and with a conserved core integration site sequence (Supplementary Figure 6), suggesting that these prophages could be horizontally transferred between different Hodors or acquired by these Hodors from closely related bacteria. Among the 147 Hodors identified from VIRE ^23^, CGVR ^24^, and metaVR ^25^, 7 were detected with prophage genomes. In addition, we *de novo* assembled 420 of the 1625 Logan samples containing Hodors (Methods), and found that 30 of these Hodor genomes harbored prophages (Supplementary Table 8). These results indicate that, although prophages are integrated in a minority of Hodor genomes, they are not rare.

### Both G1 and G2 groups of Hodors harbor a conserved, complex surface-associated gene module

We clustered the protein-coding genes encoded by all 38 curated genomes and the 1625 draft Hodor genomes based on their sequence identity. The presence and absence matrix of the protein clusters supported the definition of the two Hodor groups (Figure 2a). Both G1 and G2 Hodors encode the plasmid marker genes, including Rep_3 replication initiation protein, and ParA, ParB, and ParM/StbA involved in partitioning (Supplementary Table 2). They also encode several toxin and anti-toxin genes, and a type I-C CRISPR-Cas system with an apparently degraded repeat array (Supplementary Figure 7).

**Figure 2.**
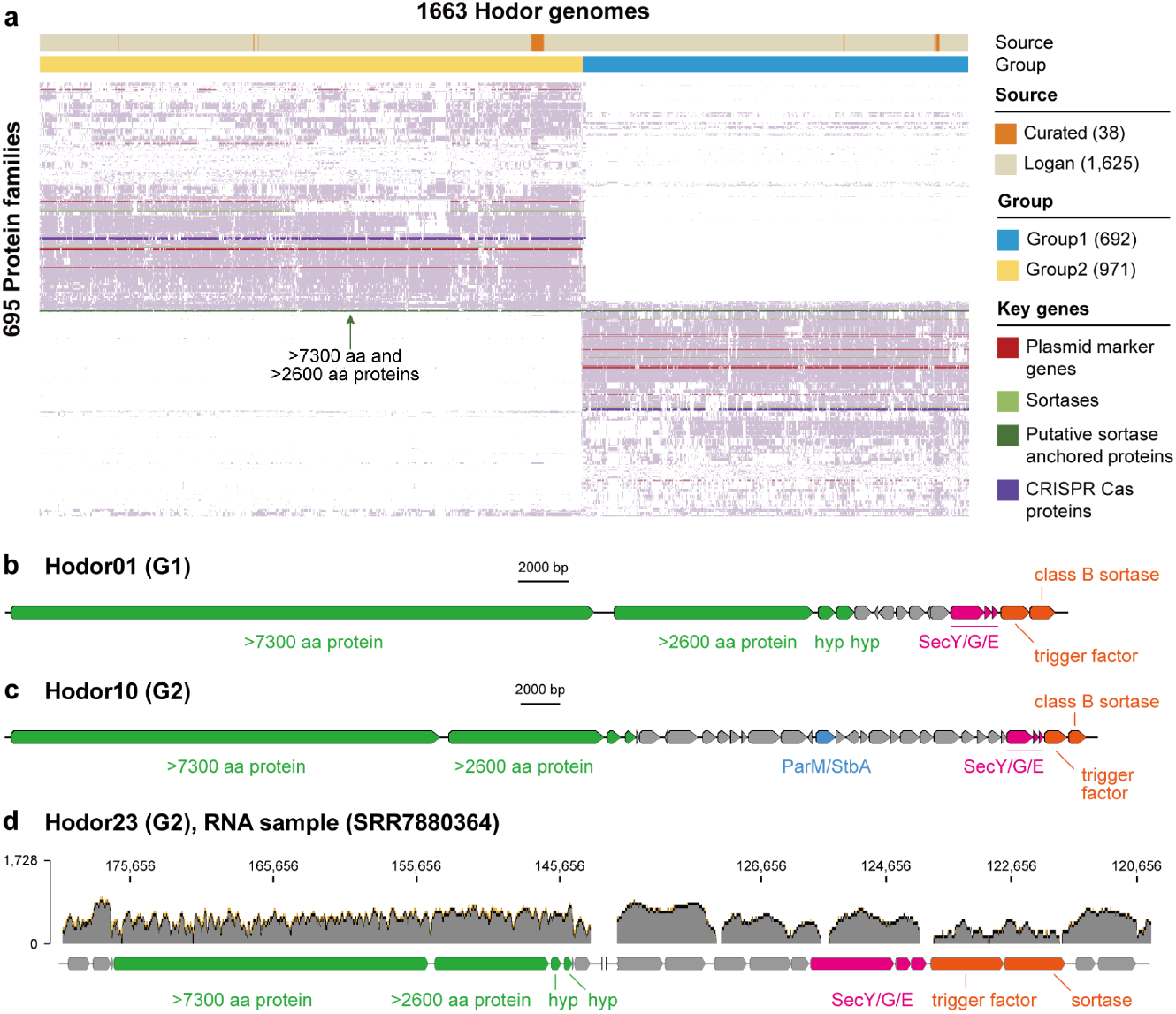
The plasmid-related genes and a conserved, complex surface-associated gene module in Hodor genomes. (a) Presence–absence heatmap of protein families across 1,663 Hodor genomes. Genome source and group assignment are indicated above. Plasmid-associated genes (including Rep_3 replication initiation proteins, ParA/ParB/ParM, and T4SS components VirD4, VirB4, and VirB6) are highlighted in red, sortase-related proteins in green, and CRISPR–Cas genes in purple. Protein families were defined by mmSeqs2 (see Methods), and only those present in at least 5% of the Hodor genomes are included here. The hierarchical clustering was performed using Euclidean distance on the binary occurrence matrix, and the clustering trees are not shown. The genomic context of surface-associated genes in (b) G1 and (c) G2 Hodor genomes. The genomes of Hodor01 of G1 and Hodor10 of G2 are used as examples to show the details of gene organization. (d) Transcriptional analysis of the surface-associated genetic module. The genome of Hodor23 from G2 is shown as an example. The DNA (SRR7880223) and RNA (SRR7880364) data were from a human stool sample (NCBI project ID, PRJNA492158). The mapping coverage is log10-scaled.

Each G1 Hodor genome encodes three sortases of classes A, B, and C, and each G2 genome encodes two sortases of classes A and B (Supplementary Table 2). In both groups, the class B sortase gene is located within a region encompassing several genes implicated in protein secretion, including a hypothetical protein (hyp), SecY, SecG, SecE, and a trigger factor (Figures 2b and c). They are among the limited number of genes that are shared by both Hodor groups (Supplementary Table 2). Other shared genes include one encoding a giant uncharacterized protein (7309-7704 amino acids; aa), and the three downstream neighbor genes, one of which encodes a protein of ∼2642 aa. These two giant proteins from G1 and G2 Hodors were assigned to the same protein clusters (Figure 2a), indicating their high conservation. Metatranscriptome analysis of a G2 Hodor genome (i.e., Hodor23) indicated that the genes for the >7300 aa protein and its three neighbor genes were co-transcribed, apparently comprising an operon (Figure 2d, Supplementary Figure 8). Two other sets of co-transcribed genes comprising putative operons near the giant protein-encoding region are the trigger factor and class B sortase genes, and the SecY, SecG, and SecE genes.

The >7300 and >2600 aa proteins harbor multiple S-layer and/or MBG (YGX type) domains that are present in a variety of bacterial extracellular proteins (Supplementary Figure 9). These proteins contain predicted signal peptides and thus are predicted to be secreted into the extracellular space, probably through a membrane pore formed by the proteins encoded by the *sec* operon. The class A and/or B sortases might position the two large proteins at the membrane, whereas the other sortases could be involved in the assembly of secreted proteins. The trigger factor likely functions as a chaperone for the giant proteins. The consistent detection of sortases and the two huge proteins predicted to be secreted with the help of the sortases suggests that the surface-associated gene module is a conserved function important for plasmid spread in both Hodor groups.

### Both G1 and G2 groups of Hodors are conjugative megaplasmids

The Hodor genomes were analyzed in detail for genes implicated in plasmid replication, partitioning, mobility, and conjugation. As described above, the replication initiation protein Rep_3 and the partitioning proteins ParA, ParB, and ParM/StbA were identified in both groups of Hodors (Supplementary Figure 4). In both G1 and G2, a large AT-rich non-coding region is adjacent to the Rep_3 replication initiation gene (Figures 3a and b), which is predicted to be the origin, OriV, for plasmid replication.

**Figure 3.**
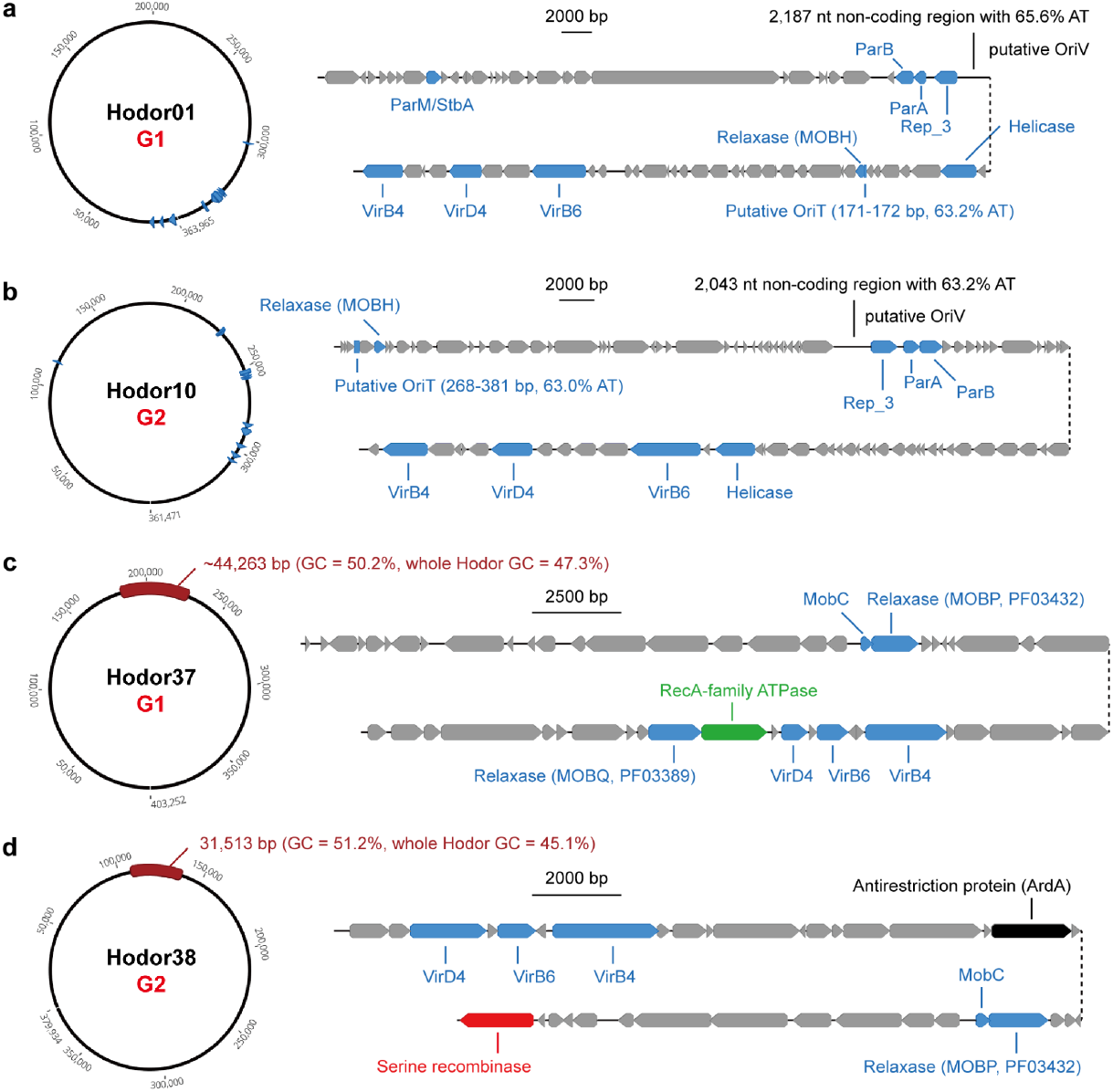
The genes related to conjugative transfer identified in Hodors. The genes identified in (a) G1 Hodors and (b) G2 Hodors. Hodor01 and Hodor10 are used as examples for G1 and G2, respectively. For both genomes, a non-coding region (putative OriV) adjacent to the gene of Rep_3 replication initiation protein is shown, with the AT content indicated. The putative OriT, which is located upstream of the MOBH-like relaxase gene, is also indicated. In Hodor01, the prophage genome is inserted into the VirD4 gene and is excluded here for display. The additional complete conjugative systems identified in (c) Hodor37 and (d) Hodor38. These two genomes were reconstructed from long-read sequencing datasets reported recently ^15^. The location of the hijacked conjugative system is indicated by the purple block, with the length and GC content shown. Note that the exact boundaries of the system in Hodor37 could not be determined precisely, which was speculated by whole genome alignment with other Hodor genomes without this system (see Supplementary Figure 12 for details). The ATPase in the hijacked conjugative system of Hodor37 belongs to the RecA family. The hijacked system in Hodor38 also encodes a serine recombinase and an antirestriction protein (ArdA).

To explore the distribution of Hodors across microbiomes, we searched 10 plasmid databases that collectively contain >852k plasmid genomes ^22^ and found only 3 distant Hodor relatives (Figure 1b, Supplementary Table 5). However, we detected 147 Hodors (excluding those with >99% identity to curated ones) and 4 distant relatives in viral databases VIRE ^23^, CGVR ^24^, and metaVR ^25^. The phylogeny built from concatenated multiple alignments of Rep_3, ParA, and ParB proteins of Hodors and the seven distant relatives suggests deep divergence of G1 and G2 Hodors (Figure 1c). We then used all 38 curated Hodor genomes individually as queries to search all the indexed NCBI metagenomic databases (all SRA made public before the end of 2023) using Pebblescout (%coverage ≥ 30; Figure 1d) and obtained 9349 non-redundant metagenomic samples containing Hodor-related reads, all from human gut or wastewater microbiomes (Supplementary Table 6). We subsequently downloaded the Logan-assembled contigs of these 9349 samples ^26^, and compared those ≥1 kbp to the curated Hodor genomes. By detecting single-copy marker genes of Hodors in the Logan assemblies (Supplementary Tables 2 and 6, Supplementary Figure 4), we obtained 1,625 draft Hodor genomes by grouping the matched Logan contigs from the corresponding samples with only one Hodor group (G1 or G2), with each genome ≥175 kbp and each contig ≥5 kbp (Supplementary Table 7). These genomes, which include 681 from Hodor G1 and 944 from Hodor G2, had an average length of 236.1 kbp, consisting on average of 18 contigs with a mean length of 13.5 kbp (Figure 1e), dramatically increasing the number of genes and genomes from both groups for more detailed downstream analyses.

Among the core type IV secretion system (T4SS) components, VirB4 is highly conserved in both groups of Hodors, with ∼48% identity between G1 and G2 (Supplementary Table 2). However, VirB6 was detected only with the help of structure prediction and transmembrane domain analysis, showing no significant sequence similarity to known VirB6 proteins (BLASTp e-value threshold = 1e-5) and low similarity between G1 and G2 Hodors (Supplementary Figure 10). VirD4, which couples the relaxosome to the T4SS, was identified in both groups of Hodors with ∼40% identity (Figures 3a and b). However, no readily identifiable relaxase-encoding gene was detected in most of the Hodor genomes. Instead, a MOB_H_-related relaxase (a member of the HD superfamily of phosphohydrolases ^27^) and a helicase encoded by a separate gene were found in both groups (Supplementary Figure 11). These two genes are located in a compact region of the Hodor genomes containing plasmid replication, partitioning, and other conjugation-related genes, including VirB4, VirD4, and VirD6. Furthermore, a non-coding region upstream of the MOB_H_-like relaxase gene was predicted to be the origin of translocation, OriT, which is essential for initiating conjugation. The putative OriT has a length of 171-172 bp for G1 and 268-381 bp for G2, with an AT content of 63.2% and 63.0% on average, respectively. Thus, Hodors encode a nearly complete conjugation machinery, including T4SS components and a MOBH-like relaxase–helicase module, consistent with a functional conjugative system.

Interestingly, one G1 (Hodor37) and one G2 (Hodor38) Hodor each harbored an additional complete conjugative apparatus (Figures 3c and d, Supplementary Figures 12 and 13). Although these two systems differed in many respects from the conjugation machinery conserved in all Hodors (in particular, in size, GC content, and gene content), they both included a relaxase of the MOB_P_ family (PF03432), a relaxome protein (MobC), coupling protein (VirD4), T4SS ATPase (VirB4), and VirB6 protein of the T4SS. We also detected an additional relaxase, belonging to the MOB_Q_ family (PF03389), associated with a RecA-family ATPase in Hodor37. Notably, the MOB_P_ family relaxase and MobC of Hodor37 and Hodor38 shared ≥96.4% and ≥95.4% identity, respectively. Further analysis confirmed that these complete conjugative systems were most similar to those in integrative conjugative elements (ICEs) of Bacillota, with the system in Hodor38 being actively horizontally transferred in the corresponding sample (Supplementary Figure 14). These results suggest that Hodors can acquire additional, autonomous conjugation modules through horizontal transfer from other mobile genetic elements (MGEs) co-existing in the same bacterial hosts.

### Hodor-associated prophages infect Bacillota and harbor targeting evolution mechanisms

The 6 prophage genomes identified in the curated Hodor genomes are 39.5-47.2 kbp in length, with a GC content of 47.6-58.9% (Supplementary Table 1). They belong to three groups defined with ≥95% identity across ≥85% of genome length (Figure 4a). We termed them prophage01, prophage02, and prophage03, respectively, with prophage01 represented by 4 genomes. Via vConTACT3 analysis, all three groups are Caudoviricetes, prophage01 and 02 belong to the same novel order but different families, with prophage03 assigned to another novel order (Supplementary Table 9). Comparison to reference phage genomes confirmed this taxonomic assignment and revealed that prophage01 and prophage02 were closely related to phages infecting Bacillota (Supplementary Figure 15). Furthermore, the UHGV-classifier assigned Bacillota as the hosts for all 6 prophages (Supplementary Table 10).

**Figure 4.**
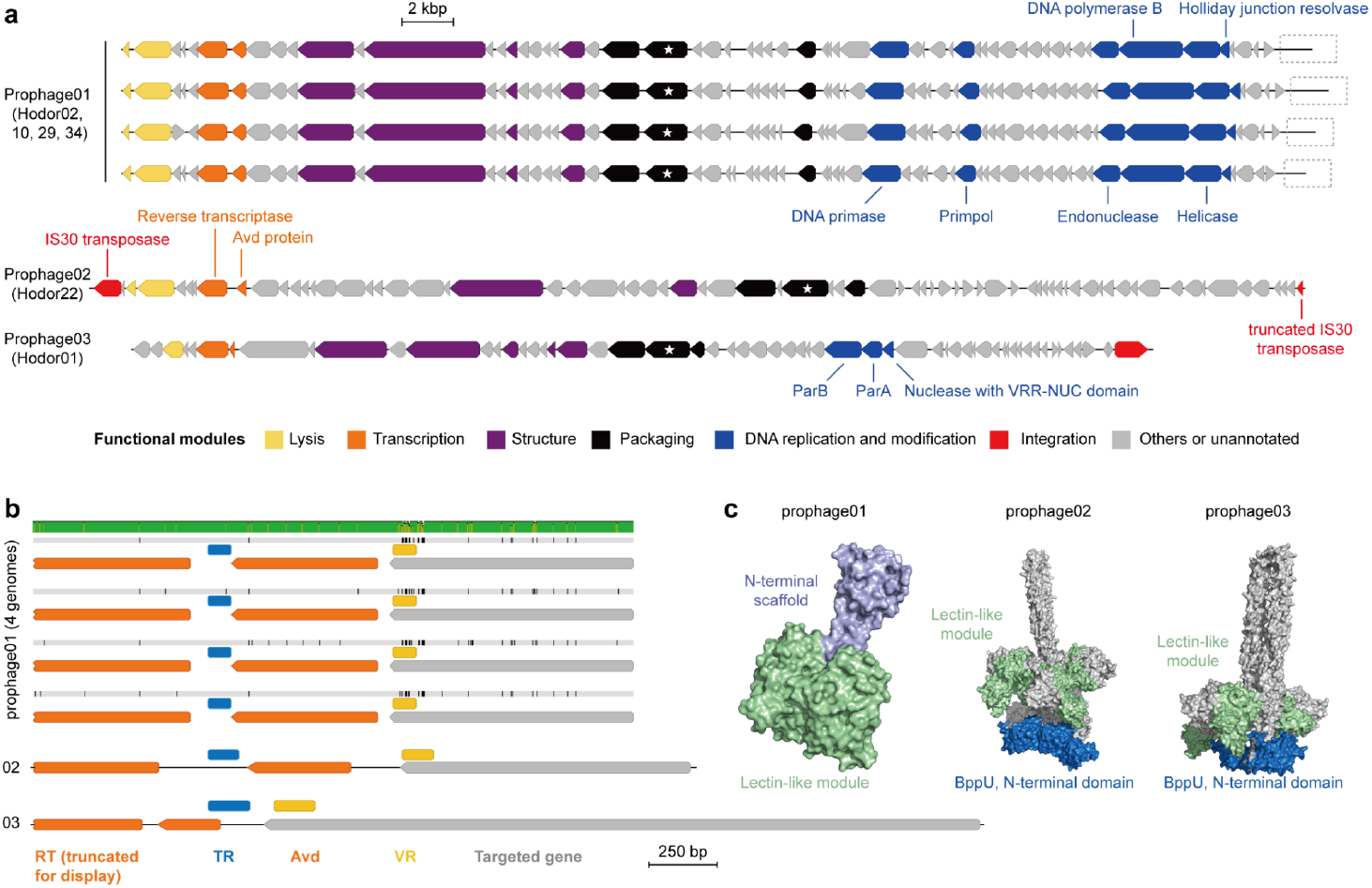
The features of Hodor-associated prophages. (a) Genomic architecture of the 6 Hodor-associated prophages in the curated Hodor genomes. The 6 prophages are assigned to three groups (01, 02, and 03) based on their overall similarity (≥95%). The genes with determined functional annotations are assigned to the corresponding module of the phage life cycle. The genes in gray are unannotated or unrelated to the listed modules. The large terminase subunit gene is indicated by a star. The dashed-line box indicates the missing prediction of the integrase gene. (b) The template region (TR), variable region (VR), and the targeted gene of the DGR systems in the Hodor-associated prophages. The four prophage genomes of the prophage01 group were aligned to indicate that the VRs are divergent among the genomes. Each black line indicates an inconsistent nucleotide among genomes. The genomes from all three prophage groups were aligned by the left boundary of their TRs. (c) Predicted protein structures of the DGR targeted genes. The structures are shown in trimers. The different modules or domains are indicated by different colors.

The prophages encode all the essential phage protein modules, including lysis, virion structure, packaging, and DNA replication (Figure 4a). Both prophage01 and prophage03 encode a serine integrase. In prophage01, the integrase gene was missed in the linear prophage genome, but can be predicted from the circular form or when integrated into the Hodor genome. Notably, the integrase in the prophages integrated in the Hodor genome is 22 aa shorter than the version from the circularized phage genome (Supplementary Figure 16). Prophage02 encodes an IS30 family transposase at the left end, with a truncated transposase at the right end, indicating that this prophage belongs to the recently identified IScream phages ^15^, in which the IS30 transposases function as integrases.

Beyond the structural and packaging modules, such as the major capsid protein and the large terminase subunit, the prophages also encode diverse DNA metabolism-related genes (Figure 4a). Prophage01 harbors a replication cassette including a DNA primase, a PrimPol-domain protein, a DNA polymerase B catalytic domain protein, and a superfamily II helicase, suggesting the potential for autonomous DNA replication following excision. Prophage03 encodes a ParAB-like partitioning module positioned between the structural genes and the integrase, as well as a VRR_NUC-domain nuclease belonging to the PD-(D/E)XK superfamily, indicative of DNA processing functions and possible maintenance during an extrachromosomal phase. In contrast, prophage02 lacks a recognizable complete replication cassette and instead carries IS30 elements at its termini, indicative of transposition-mediated mobilization. In prophage01 genomes, we additionally identified a MazF-family endoribonuclease gene. However, no canonical adjacent antitoxin was detected, leaving its functional role unresolved. The presence of these DNA replication, recombination, and maintenance modules, together with complete structural gene sets, suggests that the Hodor-associated prophages retain the genetic capacity for both lytic replication and lysogenic persistence.

We identified a reverse transcriptase (RT) gene located between the lysis and structural modules of all three prophage groups (Figure 4a). The gene upstream of the RT encodes an accessory variability determinant (Avd) protein, which is a hallmark of diversity-generating retroelements (DGRs). Structural comparison supported these annotations (Supplementary Figure 17). The presence of the DGR was corroborated using DGRscan to detect the template regions (TRs) and variable regions (VRs) (Figure 4b). The alignment of the four prophage genomes within the prophage01 group confirmed our prediction of the VRs, as the targeted genes showed high divergences in the region. The other two groups of prophages contain VRs within the 3’ region of an adjacent gene.

Although the DGR-targeted phage genes had no obvious homologs, structural predictions for the encoded proteins revealed a conserved modular architecture across prophage01–03 (Figure 4c). The targeted proteins of prophage01 genomes contain a C-type lectin-like core domain, fused to elongated N-terminal scaffolds with variable topology. In prophage02 and prophage03, the C-type lectin-like domain is further associated with an additional BppU-like N-terminal domain, suggesting these could be tail proteins, given that this domain comprises the distal tail proteins of phage TP901-1 ^28^. Despite sequence divergence between the DGR-targeted genes of prophage02 and prophage03, the overall fold and spatial organization are conserved, indicating strong functional constraints. These features are consistent with surface-exposed interaction proteins potentially involved in host recognition.

### Hodor-associated prophages are widely detected across diverse Bacillota genomes

To assess the distribution of Hodor-associated prophages in the human gut microbiome, we screened HRGM2, >20,000 assembled human gut metagenomes, and Logan assemblies (Figure 5a; Methods). In total, we identified 2351, 7861, and 219 sequences containing prophages related to prophage01, prophage02, and prophage03, respectively (Figure 5b). Protein family analysis revealed highly conserved gene content within each group, including frequent presence of DGRs. The sequences lacking DGRs were predominantly partial genomes, largely derived from Logan assemblies (Figure 5c).

**Figure 5.**
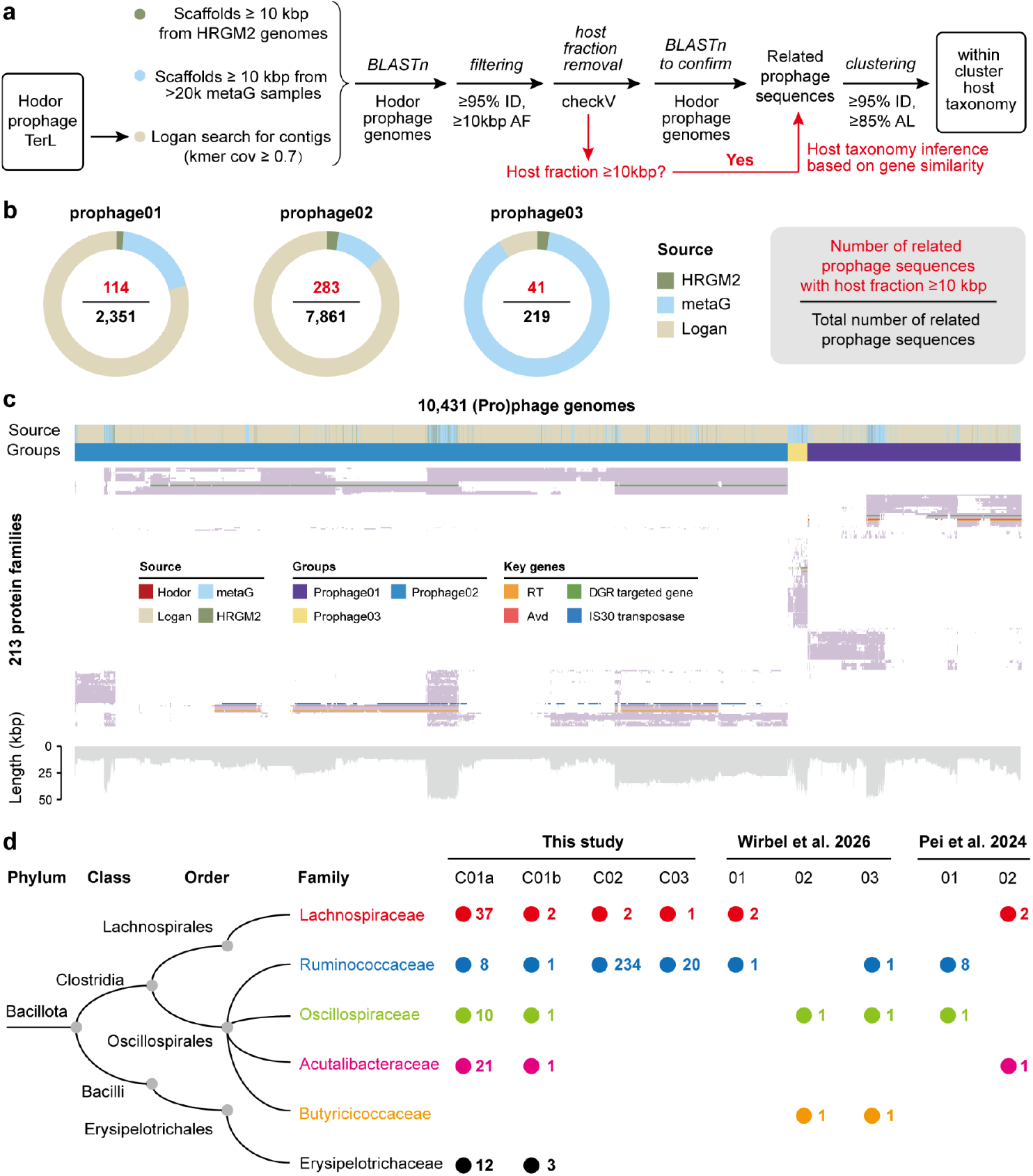
Identification of Hodor-associated prophages in bacterial genomes. (a) The analysis pipelines to identify similar prophages in HRGM2 genomes, local metagenomic assemblies (“metaG”), and Logan assemblies.(b) The number of Hodor-associated prophages identified in public data. (c) The presence/absence profiles of protein families across all the sequences related to Hodor-associated prophages. Only the protein families present in ≥5% of the sequences of an individual group are shown. The source and group assignment of the prophage-related sequences are shown at the top. Several key genes, including the reverse transcriptase, the Avd protein, the DGR targeted genes, and the IS30 transposase (prophage02 only), are highlighted. (d) The broad lysogenic host range of Hodor-associated prophages. The concept tree to the family level of the phylum Bacillota (previously Firmicutes) indicates the host assignment of the prophage members in the clusters of the corresponding prophage groups. Only the prophage clusters (4 in total) with members having bacterial hosts from at least two families are shown. The clusters were defined with ≥95% identity across ≥85% of the sequence length. The three prophage clusters reported by Wirbel et al. ^15^, and the two prophage vOTUs reported by Pei et al. ^13^, have cross-family Bacillota hosts and thus are included for comparison. For each cluster, the number of phage sequences assigned to the corresponding family is shown next to the colored circles.

Among the prophage-containing contigs, 114 (prophage01), 283 (prophage02), and 41 (prophage03) had ≥10 kb of host-derived genomic sequences, enabling confident host taxonomic assignment (Figure 5b, Supplementary Figure 18, Supplementary Table 11). Clustering of these prophage sequences at ≥95% nucleotide identity across ≥85% alignment length identified two cross-family clusters related to prophage01 (cluster01a and cluster01b), one related to prophage02 (cluster02), and one related to prophage03 (cluster03) (Figure 5d). The host ranges of cluster02 and cluster03 were dominated by bacteria of the Ruminococcaceae family, with only rare hosts from Lachnospiraceae. By contrast, the hosts assigned for cluster01a and cluster01b spanned two Bacillota classes (Clostridia and Bacilli) and multiple families, including Lachnospiraceae, Ruminococcaceae, Oscillospiraceae, and Acutalibacteraceae from Clostridia, and Erysipelotrichaceae from Bacilli. Notably, the two prophage01-related clusters showed a relatively uniform distribution across host families, consistent with a broader lysogenic host range. To determine whether the relationships among the hosts could explain the prophage genomic similarity, we examined within-cluster ANI patterns. In cluster01a, phage genomes associated with different host families were intermingled throughout the ANI dendrogram, with no clear family-specific partitioning (Supplementary Figure 19a). A finer-scale analysis of cluster02 showed a similar pattern at the genus level within Ruminococcaceae, with extensive overlap in ANI values among prophages associated with different bacterial genera (Supplementary Figure 19b).

Consistent with these findings, a recent long-read analysis by Wirbel et al. ^15^ identified three prophage clusters spanning different Bacillota families, which are among the host-families identified in this study (Figure 5d), and three out of six of the analyzed individuals studied had Hodors in their gut (Supplementary Figure 12). In addition, another previous study identified two prophage clusters in different Bacillota families, which is also in line with our observation (Figure 5d). Together, these analyses indicate that prophage genomic similarity in Hodor-associated bacterial lineages is largely uncoupled from the host taxonomy, supporting the existence of prophage populations with broad lysogenic host ranges across Bacillota.

### Multi-state coexistence of Hodor-associated prophages in the human gut

To investigate the in situ dynamics of the Hodor–prophage system, we mapped paired-end metagenomic reads to curated Hodor genomes containing integrated prophages. Read mapping revealed three co-occurring molecular populations within the same sample (Figures 6a and b), i.e., (i) Hodors lacking the prophage insertion, indicated by reads mapping exclusively to the Hodor backbone (Figure 6a, case 1), (ii) Hodors carrying an integrated prophage, supported by reads spanning the prophage–Hodor boundaries (Figure 6a, case 2), and (iii) excised circular prophage genomes, supported by reads spanning the prophage termini (Figure 6a, case 3). Thus, Hodors with and without the prophage insertion, together with circular prophage forms, can coexist within a single microbial community (see example in Supplementary Figure 2).

**Figure 6.**
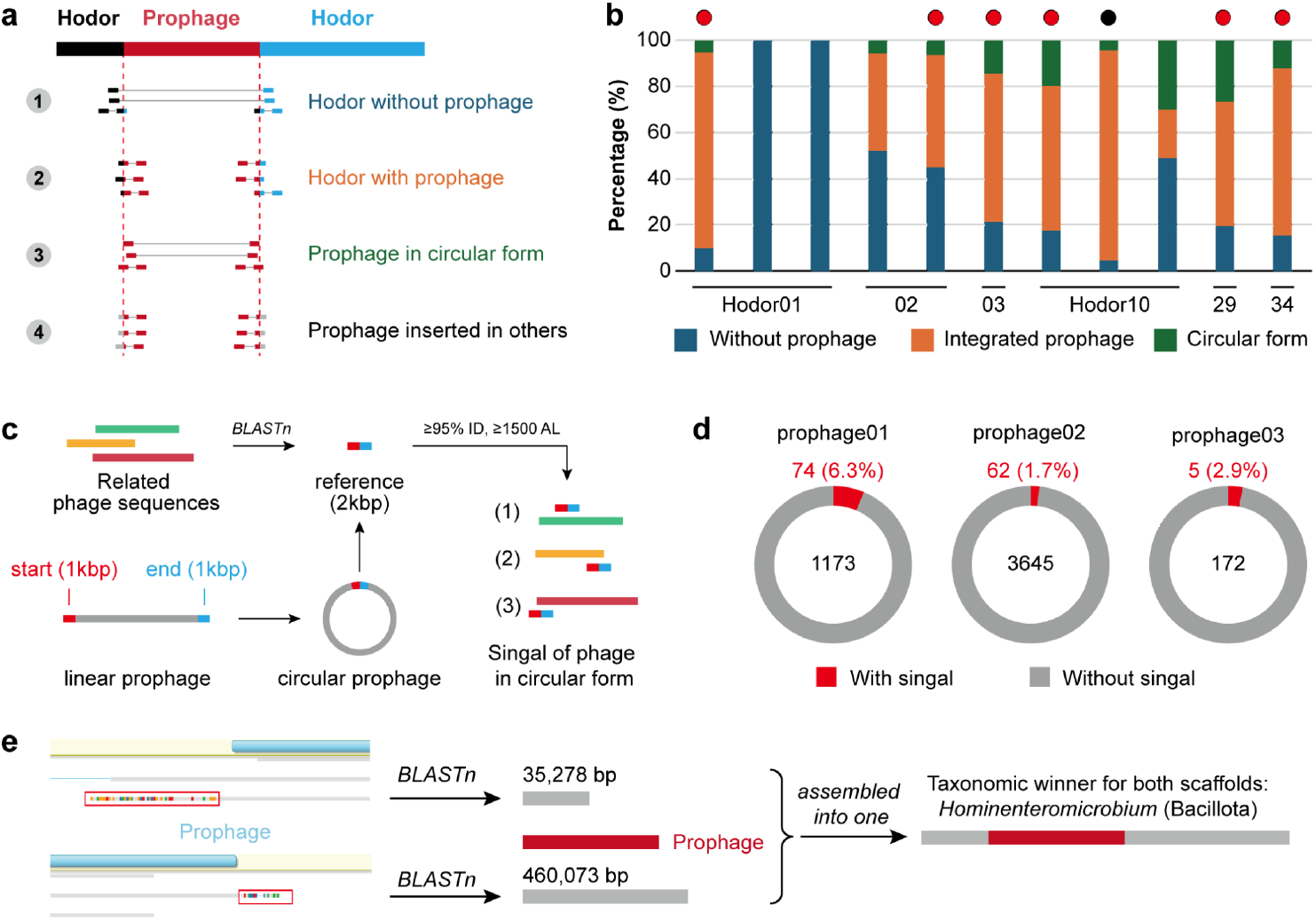
Multi-state coexistence of Hodor-associated prophages. (a) Schematic representation of the potential genetic states of Hodor and its associated prophage within a single sample. Distinct Hodor fractions are shown in different colors. (b) Distribution of detected genetic states across samples. Some Hodors were identified in multiple samples and are all included. The red circle marks the sample from which the curated Hodor genome was originally reconstructed. The black circle indicates the sample shown in panel (e). (c) Strategy for identifying excised circular prophage forms by concatenating the terminal 1 kb regions of linear prophage genomes and screening for high-identity matches spanning the artificial junction. (d) Proportion of Hodor-associated prophage sequences with or without circularization signatures. The total number of prophage-related sequences with BLASTn hits is indicated in black (center), and the subset exhibiting circular junction matches is shown in red (top). See Supplementary Figure 20 for examples supported by paired-end read mapping. (e) Example of tripartite coexistence within a single sample. In sample ERR3451245, where Hodor10 was detected at 134.4× average coverage, the same prophage is present in three states: integrated within Hodor, integrated within a bacterial chromosome, and in circular form. The red box indicates a read extending beyond the prophage boundary that did not map to Hodor. Only one from the left and one from the right boundary is shown.

To quantify the prevalence of excised forms, we concatenated the terminal 1 kb sequences of each of the six reconstructed linear prophages and screened Hodor-associated phage sequences for high-identity matches spanning the artificial junction (≥95% identity across ≥1.5 kb) (Figure 6c; Methods). Among 141 related prophage sequences retrieved from HRGM2, our metagenomic assemblies, and Logan datasets, 1.7–6.3% (3.6% on average) matched the circular junction, consistent with extrachromosomal circular genomes (Figure 6d). In multiple cases, paired-end reads directly supported the circular configuration (Supplementary Figure 20), arguing against assembly artifacts. Metatranscriptomic data further showed transcription of holin and amidase-type phage endolysin genes in a subset of samples (Supplementary Figure 21), consistent with episodic induction.

We also identified reads extending beyond the prophage boundaries that did not map to Hodor (Figure 6a, case 4), but instead aligned with independently assembled bacterial scaffolds within the same sample (Figure 6e). For example, in sample ERR3451245, Hodor10 (average depth 134.4×) co-occurred with two scaffolds (35,278 bp and 460,073 bp; comparable sequencing depth) assigned to Hominenteromicrobium (belonging to Bacillota; Figure 5d). And, these two scaffolds and the prophage could be assembled into a continuous contig, suggesting the prophage was integrated into the Hominenteromicrobium genome as well. Thus, within the same microbiome sample, the same prophage can be present (i) integrated in Hodors, (ii) integrated in a bacterial chromosome, and (iii) in the circular form.

In addition, we mapped paired-end reads from virion-enriched metagenomic datasets subjected to filtration and DNase/RNase treatment ^29^ to the prophage genomes in curated Hodor genomes (Supplementary Tables 12–14). These datasets showed broad and continuous coverage across the entire prophage genomes, including structural and packaging modules (Supplementary Figure 22), supporting the presence of virion-associated genomes in at least a subset of samples.

## Discussion

### Hodors are distinct from ICEs and phage–plasmids (P-Ps)

Although Hodors share individual features with previously described MGEs, they are mechanistically distinct from both ICEs ^30,31^ and P-Ps ^32^. ICEs are characterized by their ability to excise from and reintegrate into host chromosomes and rely on chromosomal insertion for long-term maintenance. While ICEs can mediate horizontal transfer, they are not stably maintained as autonomous replicons and are therefore not regarded as durable extrachromosomal reservoirs for complete viral genomes. In contrast, Hodors persist as large, autonomously replicating megaplasmids (Figures 3a and b) and do not depend on chromosomal integration for their maintenance. Although some Hodors occasionally acquire ICE-like conjugative modules (Figures 3c and d), these appear to be secondary insertions rather than defining features, and Hodors themselves remain stable extrachromosomal replicons.

P-Ps represent a different strategy, combining plasmid-like replication with phage-like induction and virion production ^33^. P-Ps typically lack conjugation machinery and disseminate primarily through phage-mediated infection, with their genomes maintained as intact units across life cycle transitions. Hodors differ fundamentally in that they encode complete conjugation systems and are confidently predicted to disseminate via plasmid transfer (Figure 3), while carrying prophage genomes as embedded components rather than as the principal replicative unit (Figure 1f). Thus, unlike P-Ps, Hodors decouple prophage dissemination from phage infection and instead link it to plasmid conjugation.

These distinctions position Hodors as a previously unrecognized class of composite MGEs that integrate stable plasmid maintenance, cell-to-cell transfer, and prophage carriage, providing a mechanistic framework for understanding how highly similar prophages can be distributed across phylogenetically distant bacterial lineages without invoking unusually broad infective host ranges.

### Hodors were overlooked in metagenomic datasets for various reasons

Hodors were likely overlooked in previous metagenomic surveys for several reasons. First, their overall gene content that includes virus-like components (Figure 1a) causes them to be consistently classified as viral sequences by widely used virus identification pipelines, including geNomad ^34^ and VirSorter2 ^35^, leading to their exclusion from plasmid- or host-centred analyses at early filtering stages. Second, key genetic determinants related to conjugative transfer in Hodors are difficult to identify using standard annotation approaches based on sequence similarity. For example, the T4SS component VirB6 lacks detectable homologs in curated reference databases and was only confidently annotated here through protein structure prediction combined with comparisons against multiple structural resources (Supplementary Figure 10). Furthermore, most Hodors lack the most common relaxases, such as those of the MOBF family, encoding an HD nuclease (MOBH family relaxase) and a helicase in separate genes instead (Supplementary Figure 11). Third, Hodor genomes that carry prophages are inherently challenging to recover from short-read assemblies ^36^. Phage genomes can coexist in multiple contexts, including bacterial chromosomes and/or as free viral particles, preventing co-assembly of plasmid and prophage sequences into a single sequence if the prophage is present in only a small subset of the plasmid (Figures 1 and 6, Supplementary Table 1). Although long-read sequencing offers an opportunity to recover longer and more complete Hodor or Hodor-like genomes (Supplementary Figure 12), its full potential will likely be realized only when combined with short-read mapping to resolve heterogeneous population states of the associated prophages (Supplementary Figure 14). Together, these factors explain why Hodors have escaped detection in previous metagenomic datasets and highlight the importance of integrative analytical strategies for uncovering such composite and potentially ecologically impactful mobile elements.

### Ecological distribution and evolutionary features of Hodors

Beyond their role in prophage carriage, Hodors exhibit ecological and evolutionary features that distinguish them from previously described MGEs ^37^. Although they form two deeply diverged groups of megaplasmids, Hodors share highly conserved gene modules, including, apart from the closely related prophages, a large gene module encoding surface-associated proteins, in particular, two giant proteins (>7300 aa and >2600 aa), predicted to be secreted, and likely anchored on the cell surface via Hodor-encoded sortases ^38^. The conservation of these features across divergent Hodor groups suggests that all Hodors occupy related ecological niches and are hosted by bacteria sharing similar cell envelope architectures. Consistently, most Hodor-encoded genes show the highest similarity to homologs from Bacillota, indicating lineage-level constraints on plasmid persistence and transfer.

Within individual Hodor groups, distinct prophages associate with closely related plasmid backbones, and conversely, closely related prophages are found in divergent Hodors from different groups (Figure 1f), indicating horizontal mobility of prophage among Hodors and/or between Hodors and host bacteria rather than strict co-evolution. Moreover, a subset of Hodors carry complete ICE-like conjugative systems (Figures 3c and d). This observation suggests two non-exclusive evolutionary scenarios: Hodors might act as vehicles for ICE dissemination, or alternatively, might hijack ICE-encoded conjugation systems to enhance their own mobility ^33^. Such interactions imply that Hodors participate in dynamic networks of MGEs, blurring boundaries between plasmids and other integrative elements.

From an evolutionary perspective, Hodors combine deep sequence divergence with strong functional constraint. Core replication and partitioning modules are moderately conserved between groups (Figures 3a and b), whereas relaxase-related components exhibit greater sequence divergence yet retain structural similarity (Supplementary Figure 11), consistent with function conservation. Together, these features position Hodors as adaptable, composite elements embedded within broader exchange networks of MGEs and serving as vectors for prophages and possibly other MGEs.

### Temperate phage dissemination mediated by conjugative megaplasmids

Our data support a model in which conjugative megaplasmids facilitate cross-lineage dissemination of temperate phage genomes across Bacillota. Despite plasmids being well known for carrying diverse small MGEs ^39^, such as transposable elements, reports of complete prophage-harbouring plasmids are scarce ^40,41^. In our data, we found a subset of Hodors carrying complete, intact prophages (Figure 1) that are highly similar despite being associated with bacterial hosts spanning distinct Bacillota families and, in some cases, different classes (Figure 5). Within cross-family clusters, prophage genomes show extensive overall similarity and are intermingled across host taxonomic groups (Supplementary Figure 19), consistent with recent or recurrent movement across phylogenetic boundaries rather than strict host-linked divergence.

In this scenario, prophages are disseminated as embedded components of conjugative plasmids and subsequently integrate into recipient chromosomes following plasmid transfer. Conjugation thus provides a route for temperate phage genomes to enter new host lineages without requiring immediate productive infection, effectively decoupling lysogenic host range from infective host range. This mechanism offers a plausible explanation for the widespread occurrence of closely related prophages across phylogenetically distant bacteria in the human gut ^13–15^.

Notably, all three groups of Hodor-associated prophages encompass DGRs, and the predicted variable regions map to proteins with conserved, tail-associated architectures, including C-type lectin-like domains (Figure 4c). Such DGR-linked, structurally constrained variability is consistent with ongoing selection on host-interaction functions ^42,43^. We therefore propose that plasmid-mediated transfer and infection-mediated spread are complementary rather than mutually exclusive. Hodors could enable cross-lineage delivery and chromosomal establishment of prophage genomes, whereas DGR-driven diversification could facilitate subsequent tuning of host-recognition determinants, enabling formation of phage particles upon prophage induction, egress, and productive infection in some of the hosts. The detection of circular forms (Figure 6) and virion-associated reads (Supplementary Figure 22), together with transcriptional signatures of lysis-related genes in a subset of samples (Supplementary Figure 21), supports the potential for lytic amplification of Hodor-associated prophages once compatibility is achieved.

Together, these findings suggest an expanded conceptual framework for temperate phage ecology ^44^. Hodors represent composite mobile genetic platforms that integrate stable plasmid maintenance, cell-to-cell transfer, and prophage carriage, thereby potentially enabling cross-lineage movement of viral genomes. By coupling conjugative dissemination with adaptive diversification of host-recognition modules, this system reconciles the apparent paradox of narrowly infective phages whose prophages nonetheless display broad phylogenetic distributions. Conjugative megaplasmids thus might constitute an underappreciated driver of viral persistence, diversification, and host-range evolution in complex microbial ecosystems. Experimental validation will be essential to determine the frequency and ecological impact of this plasmid-mediated dissemination route relative to infection-mediated spread.

## Conclusion

This study uncovers conjugative megaplasmids that embed complete, intact temperate phage genomes, revealing a class of composite mobile elements that expand current views of phage dissemination in complex microbiomes. By enabling prophages to overcome host-range barriers and integrate into diverse bacterial genomes, Hodors suggest a plasmid-mediated route for cross-lineage viral spread. Future work should focus on experimental validation of Hodor-mediated transfer, chromosomal integration dynamics, and induction competence, as well as direct tests of host range and DGR-driven adaptation of phage proteins involved in host recognition. Systematic identification and experimental characterization of Hodor-like elements across environments will be essential to quantify the ecological impact of plasmid-mediated phage dissemination. Such efforts will help establish a mechanistic framework for understanding viral dispersal, persistence strategies, and host-range evolution in microbial ecosystems.

## Methods

### Identification of Hodor megaplasmids

In our analysis of the huge phage genome collection (HPGC) ^16^, two groups of genomes caught our attention because they encompass genes for sortases, which have not been reported in phages previously. Sortases are transpeptidases that covalently anchor surface proteins to the cell envelope in Gram-positive bacteria ^20^. These two groups of genomes were defined at ≥90% nucleotide identity and across ≥80% genome length, and contained 9 and 10 genomes each group, respectively. We defined them as group1 (G1) and group2 (G2). All of the genomes were originally reconstructed from human gut metagenomic samples ^17–19^. Only two genomes in each group encoded a large terminase (*TerL*), a conserved, hallmark gene present in all tailed phages, that was identified using sequence and structural searches, although paired-end reads mapping indicated that some genomes lacking TerL genes are circular.

### Manual genome curation

To curate the Hodor genomes, the corresponding paired-end read files were retrieved via Pebblescout ^21^ online search with default parameters. The corresponding paired-end reads of each draft Hodor genome were determined if the Pebblescout “%coverage” was >99. The paired-end reads were downloaded from NCBI and assembled using metaSPAdes version 3.15.2 ^45^ with the kmer set of “21,33,55,77,99” (when read length is 100 bp) or “21,33,55,77,99,127” (when read length is 150 bp). Among the 19 draft Hodor genomes, 17 had paired-end reads available. The *de novo* assembled scaffolds were then searched against the corresponding draft Hodor genome using BLASTn for related scaffold(s). Then the paired-end reads were mapped to the identified scaffold(s) to identify and fix any assembled errors or gaps (i.e., Ns in scaffolds), and also to extend the scaffolds to a circular genome if possible, as previously described ^46^. The details are available at https://ggkbase-help.berkeley.edu/genome_curation/scaffold-extension-and-gap-closing/. The scaffold extension is based on the unplaced paired-end reads, which are, for example, within paired reads of read_1 and read_2. If read_1 is mapped to the reference and read_2 is not, then read_2 is called the unplaced paired-end read. The tool of shrinksam (available at https://github.com/bcthomas/shrinksam) was previously developed to output both reads when only one read is well mapped, without saving paired reads that are not mapped. Once a Hodor genome was manually curated to a circular genome (that is, with paired-end reads spanning the two ends of the scaffold) or could no longer be extended any more, the reads were mapped to the scaffold, allowing no mismatch, to confirm that no more errors exist.

### Identification and curation of additional Hodor genomes

To obtain more high-quality Hodor genomes with inserted prophage or with metatranscriptomic data available, we searched Pebblescout ^21^ for the 19 Hodor genomes from HPGC (Supplementary Table 1). The output tables were manually checked for metagenomic samples with paired metatranscriptomic datasets, and identified at least several from three NCBI projects, i.e., PRJNA492158, PRJNA354235 ^47^, and PRJNA818303 ^48^. The metagenomic reads were downloaded and *de novo* assembled. And we also randomly picked and assembled the paired-end assembled around 100 of the identified metagenomic samples with a Pebblescout “%coverage” of >70. All the paired-end reads were quality controlled using fastp with default parameters ^49^, and the clean reads were assembled using metaSPAdes as described above.

The short reads and long sequencing reads from a recent study ^15^ were re-analyzed for Hodor genomes. Long reads were quality-controlled using fastplong version 0.3.0 with default parameters ^50^. The cleaned long reads were assembled using metaFlye version 2.9.6-b1802 using default settings ^51^. The assembled contigs were compared against the curated Hodor genomes from HPGC using BLASTn. Any contigs with >90% sequence identity over an aligned length exceeding 175 kb were retained for further manual genome curation. The manual curation and extension of additional Hodor genomes were the same as described earlier, except that the search for target scaffold(s) for curation was based on a BLASTn search against the HPGC Hodor genomes.

### Protein-coding gene prediction, protein structure prediction, and analysis

The protein-coding genes from the 38 curated Hodor genomes were predicted using Prodigal version 2.6.3 (-m -p meta) ^52^. The predicted protein sequences were clustered with ≥70% identity across ≥80% length of the shorter ones using mmSeqs2 version 15.6f452 (--min-seq-id 0.7 -c 0.8 --cov-mode 1 --cluster-mode 2) ^53^. The clustering output was transferred into a matrix (Supplementary Table 2), and the protein clusters in at least three genomes were analyzed using hierarchical clustering based on Euclidean distance (Supplementary Figure 4).

Protein structures of the protein cluster representatives (1290 in total) were predicted using ColabFold ^54^ with the parameters ‘--num-recycle 3, --use-gpu-relax, --amber, and --stop-at-score 70’. For each protein, only the rank_001 model with an average predicted local distance difference test (pLDDT) score ≥ 70 was retained for functional annotation. The structures were searched against the Big Fantastic Virus Database (BFVD) ^55^, the Protein Data Bank (PDB) ^55,56^, and the NCBI Conserved Domain Database (NCBIfam) ^57^, using Foldseek version 9.427df8a with the easy-search workflow (-c 0.7 --cov-mode 0) ^58^. Only those with an alntmscore ≥ 0.5 and e-value < 1e-5 were thought to be reliable matches.

We also predicted the functional domains of the protein cluster representatives using HMMsearch from HMMER version 3.3.2 ^59^ against the Pfam HMM database ^60^. The output file was parsed using cath-resolve-hits version 0.16.10-0-g99edb28 ^61^. The domains with an indp-evalue of <1e-5 were thought to be reliable.

Some of the key genes described in the Results are manually confirmed based on both protein structure comparison and predicted functional domains. All these results are summarized and included in Supplementary Table 2 or displayed in the corresponding figures and tables.

### Identification and confirmation of prophage genomes in Hodor genomes

The complete, intact prophage genomes in the Hodor genomes were determined and confirmed by the annotation of protein-coding genes in the corresponding regions. The exact insertion boundaries of the prophage genomes were determined by mapping the paired-end reads to the curated Hodor genomes. There are several types of paired-end reads mapping information that provide information that are useful in the determination (see Supplementary Figure 2 for examples), including (1) junction reads, (2) paired-end reads spanning the two ends of the prophage region, and (3) paired-end reads mapped to the Hodor genomes while spanning the prophage region.

### Analysis of Hodor genes similar to those from Bacillota

We compared the protein-coding genes encoded by all 38 curated Hodor genomes (with those from prophages excluded) against the reference proteins in the UniProtKB database ^62^ using MMseqs2 version 15.6f452 ^53^. The taxonomy information of each Hodor protein was based on the first UniProtKB hit. To evaluate the relatedness of the Hodor proteins most similar to those from Bacillota, phylogenetic analyses of several proteins were performed, including sortase B and the Sec translocon components SecY, SecG, and SecE. Protein sequences from Hodors were first searched against the HRGMv2_100 protein database ^63^ using BLASTP, with no E-value cutoff applied and a maximum of 2,000 target sequences retained per query. The resulting HRGMv2_100 protein sequences were searched against the Pfam database using hmmsearch, and only those assigned to the same Pfam family as the corresponding Hodor gene were retained. The filtered HRGMv2_100 protein sequences were then clustered using CD-HIT version 4.8.1 ^64^ with parameters ‘-c 0.9 -aS 0.8 -G 0’. The representative HRGMv2_100 protein sequences and the Hodor protein sequences were aligned using MUSCLE version 5.3.linux64 [d9725ac] ^65^. Poorly aligned positions were trimmed with TrimAL version v1.4.rev15 with the parameter set as -gt = 0.2 ^66^. Maximum-likelihood phylogenetic trees were inferred using IQ-TREE version 2.4.0 (-m LG+G4 -bb 1000) ^67^.

### Identification of Hodor and relatives in public databases

To reveal the distribution of the two groups of Hodors and their relatives, we first searched the 10 plasmid databases in PlasmidScope ^22^, which contain the plasmids from PLSDB ^68^, COMPASS ^69^, mMGE ^70^, IMG/PR ^71^, TPA ^72^, and those from ENA ^73^, Kraken2 ^74^, GenBank ^75^, DDBJ ^76^, and RefSeq ^77^, and collectively have over 852,600 plasmid genomes. The plasmid genomes from all these databases were downloaded from https://plasmid.deepomics.org/, and protein-coding genes were predicted using Prodigal version 2.6.3 (-m -p meta) ^52^. Then the predicted protein sequences were searched against the sequences of Rep_3 replication initiation protein, ParA, ParB, and ParM/StbA from the 38 curated Hodor genomes. Once a query plasmid genome has at least one protein with ≥50% identity to any of the four Hodor proteins, the plasmid genome will be retained for further analysis. If a retrieved plasmid genome shared ≥90% similarity with ≥10 kbp genomic length to any of the 38 curated Hodor genomes, it was assigned as a Hodor genome and was subsequently clustered with curated Hodor genomes for group assignment; otherwise, it was assigned as a distant relative of Hodors. The same analysis was conducted for Hodors and their relatives in public viral databases, including VIRE ^23^, CGVR ^24^, and metaVR ^25^. To reveal the phylogenetic relatedness of the two groups of Hodors, a phylogenetic tree was built using the concatenated protein sequences of Rep_3, ParA, and ParB (in this order) from all curated Hodor genomes and their distant relatives. The identified Rep_3, ParA, and ParB were respectively aligned using MUSCLE version 5.3.linux64 [d9725ac] with default parameters ^65^; the alignments were concatenated and filtered to remove any columns comprising ≥90% gaps using TrimAL version v1.4.rev15 ^66^. The tree was constructed using IQ-TREE version 2.4.0 with “-bb 1000” and “-m LG+G4” ^67^.

### Reconstruction and analysis of draft Hodor genomes

To obtain more Hodor genomes for further analyses and to reveal their distribution as well, we searched all 38 curated Hodor genomes using Pebblescout ^21^ against the “Metagenomic, Volume 1” and “Metagenomic, Volume 2” databases. The hit SRA samples with a minimum Coverage% of 30 were retained. All the retained hit SRA samples from all curated Hodor genomes were duplicated to obtain a non-redundant sample list (Supplementary Table 6). The Logan assembled contigs ^26^ of these samples were downloaded and filtered to keep those with a minimum length of 1000 bp. The Logan contigs ≥ 1000 bp were then searched against the 38 curated Hodor genomes using BLASTn. The contigs with a minimum similarity of 90% with the curated Hodor genomes across at least 70% of contig length were considered as hits. We also predicted protein-coding genes from the Logan contigs ≥1000 bp using Prodigal version 2.6.3 (-m -p meta) ^52^, and compared the predicted proteins against the Rep_3 replication initiation protein, ParA, ParB, and ParM/StbA of the two groups of Hodor using BLASTp, allowing ≥90% identity. If a given sample had only one type of Hodor, which was based on the BLASTp results, then the BLASTn hits from that sample were grouped together to represent the draft Hodor genome. Only those draft Hodor genomes with a collective contig length of 175 kbp were retained for further analyses. Here, a genome with at least 175 kbp represents ≥50% identity, given that the complete Hodor genomes have an average length of approximately 351 kbp.

### Protein family analysis

Protein family analyses were performed for three sets of genomes in this study, including (1) the 38 curated Hodor genomes, (2) the 38 curated Hodor genomes along with the 1625 Logan-retrieved draft Hodor genomes, and (3) the 10,431 Hodor-associated prophage genomes. The protein-coding genes were predicted using Prodigal version 2.6.3 (-m -p meta) ^52^, then clustered into protein families using mmSeqs2 version 15.6f452 (“--min-seq-id 0.7 -c 0.8 --cov-mode 1 --cluster-mode 2”) ^53^. For each set of genomes, the presence/absence matrix of the defined protein families was hierarchically clustered based on Euclidean distance.

### Architectural analysis of two putative sortase-anchored surface proteins

Sortase-anchored surface proteins typically exhibit a conserved architectural paradigm comprising an N-terminal signal peptide for secretion and a C-terminal cell wall sorting signal (CWSS) that includes a sorting motif, a hydrophobic transmembrane (TM) helix, and a positively charged tail. Across 38 Hodor genomes, we consistently identified two large proteins (>7300 aa and >2600 aa) that conform to this paradigm. Signal peptides were predicted using DeepTMHMM v1.0 (https://services.healthtech.dtu.dk/services/DeepTMHMM-1.0/). The TM helix in the CWSS was identified as a ∼15–25-aa continuous hydrophobic segment (rich in I/L/V/F/M/A). The positively charged tail was defined as the K/R-enriched residues immediately following the TM helix. Candidate sorting motifs were annotated from the upstream of the TM helix based on conserved sequence patterns within multiple alignments using MUSCLE ^65^. To further characterize the functional region, the consensus sequences were searched against the NCBI nr database using online BLASTp, and conserved-domain annotations were extracted from the corresponding graphic summaries.

### Analysis of HD nuclease relaxase and helicase genes

We did not identify any canonical relaxase-encoding genes in most curated Hodor genomes. Instead, we detected a short MOBH-like relaxase (HD nuclease) gene and a helicase gene in both G1 and G2 Hodors, supported by structural similarity to a TIGR03760 representative structure ^57^ and by HHsearch ^78^ hits to the TIGR03760 HMM. These genes were consistently located adjacent to the plasmid replication, partitioning, and conjugation-associated T4SS genes. To reveal the conservation of the key catalytic residues of the HD nucleases, we first recruited homologs through a PSI-BLAST search with 1 round, with the cutoffs being E-values < 1E-20 and coverage >92.5%. Then, we produced multiple sequence alignments using MUSCLE (version 5.3) ^65^ and produced sequence logos using the genotypst Typst package (https://github.com/apcamargo/genotypst). To reveal the phylogenetic relatedness of the HD nucleases with MOBH relaxases, we retrieved close homologs using BLASTp. Then a phylogeny was reconstructed from the Hodor HD nucleases, the retrieved proteins, and the known MOBH relaxases using IQ-TREE (version 3.0.1) ^79^. The functional domains of the Hodor helicases were predicted using InterProScan (https://www.ebi.ac.uk/interpro/search/sequence/), and the diagram was made based on the prediction results. The structures of Hodor HD nucleases and helicases were predicted using ColabFold ^54^ with the parameters ‘--num-recycle 3, --use-gpu-relax, --amber, and --stop-at-score 70’, and visualized and aligned in ChimeraX version 1.11 ^80^.

### OriV and OriT prediction

In plasmids, the origin of vegetative replication (OriV) is typically located adjacent to replication initiation protein–encoding genes. In all curated Hodor genomes from both groups, we identified a non-coding region exceeding 2000 nt immediately upstream of the Rep_3 genes, which was therefore considered a putative OriV region based on genomic context. The putative OriT regions were identified based on their proximity to the HD nuclease (MOBH-family relaxase) genes. In G1 Hodors, a non-coding region of 171–172 bp was consistently detected immediately upstream of the HD nuclease gene. In G2 Hodors, a conserved non-redundant region ranging from 268 to 381 bp was identified upstream of the HD nuclease gene, separated by a hypothetical protein-coding sequence. The secondary structure folding of the candidate OriV and OriT regions was performed using UNAFold (https://www.unafold.org/) under default DNA folding parameters to evaluate the presence of stable stem–loop structures. The AT content of the predicted OriV and OriT regions was calculated and compared with that of the complete Hodor genomes. As these predictions were based solely on genomic context and computational analyses, the exact boundaries and functional validation of OriV and OriT could not be determined.

### Metatranscriptomic analyses of Hodors

Paired metatranscriptomic data are available for three of the G2 curated Hodor genomes (Hodor23, Hodor35, and Hodor36), which made it possible to evaluate their transcriptional activities. In detail, three RNA samples were for Hodor26, two for Hodor35, and one for Hodor23. The RNA data were downloaded and quality controlled using fastp with default parameters. The quality RNA reads were mapped to the corresponding curated Hodor genomes using Bowtie2 version 2.5.4 ^81^. The Shriksam tool (https://github.com/bcthomas/shrinksam) was applied to exclude unmapped pairs from the SAM file, and was subsequently sorted and converted to the BAM format using SAMtools ^82^. The obtained BAM format file was imported into Geneious Prime Build 2025-03-24 ^83^ for visualization and analysis. The RNA reads were remapped in Geneious Prime using the “Map to Reference(s)” function, allowing no mismatch. The resulting RNA reads mapping profiles were shown accordingly.

### Identification of hijacked conjugative systems in Hodor genomes

The hijacked conjugative systems were first identified in Hodor37 (G1) and Hodor38 (G2) via comparing their genomes against other curated Hodor genomes from the same group (Supplementary Figure 12). In each of these two genomes, we found a region that was absent in other curated Hodor genomes, and conjugation-related genes were encoded by the region. Long reads mapping confirmed that the presence of the additional conjugative systems in these two genomes was not due to assembly errors (Supplementary Figure 13). Comparison the encoding genes in the region of Hodor37 via online BLASTp, and mapping the paired-end short reads to the curated Hodor38 genome, suggested that these systems were from integrative conjugative elements (ICEs) (Supplementary Figure 14). Notably, the additional conjugative system identified in Hodor38 was also identified in a bacterial genome in the same sample, and the corresponding bacteria had a lower relative abundance than Hodor38 (ratio was ∼1:3 according to the sequencing coverage calculated using paired-end short reads).

### Identification of Hodor-associated prophage genomes in public databases

To assess the distribution of Hodor-associated prophages in bacterial genomes and microbiomes, we searched three resources, i.e., HRGM2, >20,000 human gut metagenomic assemblies (hereafter “metaG”, which were assembled for other research projects), and Logan assembled contigs ^26^. The 6 prophage genomes identified from curated Hodor genomes were used as references (Supplementary Table 1). For HRGM2 and metaG scaffolds, BLASTn searches were performed using the 6 Hodor-associated prophage genomes as references. Scaffolds sharing ≥95% nucleotide identity over ≥10 kbp were considered candidate matches. For Logan contigs, we first searched the first 1000 bp (the N-terminal) of the large terminase subunit genes from the 6 prophages against the Logan database (https://logan-search.org/dashboard). Contigs from samples with k-mer coverage ≥0.7 were retrieved and filtered to retain sequences ≥1000 bp. These contigs were subsequently searched against the 6 prophage genomes using BLASTn, and sequences with ≥95% identity across ≥10 kbp were considered matches. All matched scaffolds or contigs were analyzed using CheckV ^84^ version 1.0.3 to separate viral and host fractions where applicable. The viral fractions were re-aligned to the 6 reference prophage genomes using BLASTn (≥95% identity across ≥10 kbp) to confirm their assignment. The samples having these contigs are listed in Supplementary Tables 12–14.

For sequences containing host-associated regions ≥10 kbp, host taxonomy was inferred. Protein-coding genes were predicted from the host fraction using Prodigal v2.6.3 (-m -p meta) ^52^ and searched against the UniProtKB database using MMseqs2 v15.6f452 (--search-type 1 --orf-filter-e 1e-4 --max-seqs 1) ^53^. The MMseqs2 UniProtKB database ^62^ was built and linked to NCBI taxonomy, and taxonomic annotations were extracted directly from the best hits. Host taxonomy was assigned using a last common ancestor (LCA) approach: if ≥50% of genes were assigned to the same genus, that genus was reported; otherwise, the assignment was determined at the family, order, class, or phylum level using the same ≥50% criterion. All predicted hosts were classified within Bacillota. Because NCBI taxonomy is not fully synchronized with the updated GTDB framework, we constructed an independent validation database. One representative genome from each Bacillota genus in GTDB ^85^ (v226) was randomly selected, and protein-coding genes were predicted using Prodigal (-m -p meta) ^52^. Taxonomic information was embedded in protein headers, and a BLASTp database (Bacillota_proteins) was constructed. Host-derived proteins were searched against this database using BLASTp (e-value ≤1e-5), and taxonomy was reassigned using the same LCA strategy. To validate this approach, we leveraged the fact that HRGM2 is a genome-resolved dataset: each prophage-matching scaffold is associated with a specific HRGM2 genome. We therefore compared our inferred host taxonomy (from the host fractions) against the GTDB-Tk taxonomy (version 2.4.1) ^86^ of the corresponding HRGM2 genomes and found complete agreement (Supplementary Table 11). The full workflow is illustrated in Supplementary Figure 18.

### The taxonomy and potential hosts of Hodor-associated prophages

To reveal the taxonomy and potential bacterial hosts of the prophage identified in Hodor genomes, we submitted the prophage genomes to viptree (https://www.genome.jp/viptree/) ^87^, which uses protein families from both uploaded phages and dsDNA phages for inference. The taxonomy and potential hosts of Hodor-associated prophages were accordingly predicted based on the most closely related reference phages.

### Modular architecture analysis of DGR-associated C-type lectin-like proteins

To characterize the modular organization and potential functional domains of typical DGR-associated C-type lectin-like proteins, a hierarchical annotation strategy combining sequence- and structure-based approaches was applied. Protein sequences were analyzed using InterProScan version 5.62–86.0 to identify Pfam, InterPro, or other conserved domains. AlphaFold2 was used to predict the three-dimensional structures of the target proteins. Given that tail-associated host-recognition proteins commonly assemble as oligomers, structural predictions were performed using AlphaFold-Multimer. For each target protein, three identical amino acid sequences were provided to generate trimeric models, and the highest-confidence trimer structure was selected for subsequent structural analyses.

To assess structural homology independent of sequence similarity, each predicted structure was compared against the PDB using DALI to evaluate potential folds and modular features. Merizo version 1.0 was applied to partition each protein into discrete folding modules, which were then cross-referenced with InterPro/Pfam annotations: modules with clear sequence-based domains (e.g., BppU_N) were labeled accordingly, while modules lacking sequence annotation were labeled based on structural features alone. All modules were visualized and annotated in PyMOL version 3.0.3 (Schrödinger, LLC), with N-terminal scaffolds and lectin-like modules distinguished by color.

### Circular signal of Hodor-associated prophages

To detect excised circular forms independent of integration context, we concatenated the start 1000 bp and end 1000 bp of the complete linear prophage genomes that we reconstructed from Hodor genomes as a reference for comparison. The retrieved phage sequences related to Hodor-associated prophages were searched against this 2000 bp fragment, and those with a region having over 95% similarity across over 1500 bp were considered as being phages in circular form.

## Data availability

The 38 curated Hodor genomes and the 6 prophage genomes in curated Hodor genomes are available at Figshare (https://doi.org/10.6084/m9.figshare.31385365).

## Acknowledgments

We thank A. Murat Eren for the insightful discussion in the initial stage. We thank Dr. Jillian F. Banfield for commenting on the manuscript. We thank Junhyeong Kim for providing helpful suggestions on folding the SecYGE complexes. We thank the Supercomputing Center at the University of Science and Technology of China for its support of ColabFold analyses. L-X.C. was supported by the Research Program of the University of Science and Technology of China KY2400000036.

## Author contributions

L.Y., A.P.C., E.V.K., and L-X.C. designed the study. L.Y. and L-X.C. performed the metagenomic assemblies and curated the genomes. Y.Q., A.P.C., E.V.K., and L-X.C. analyzed the plasmid marker genes. L.Y., J.W-R, and L-X.C. performed the analyses of the surface-associated gene module. L.Y. and L-X.C. reconstructed the draft Hodor genomes from Logan assemblies and performed all related analyses. Y.Q., A.P.C., and L-X.C. performed the DGRs-related analysis. L.Y. and L-X.C. mined the Hodor-associated prophage sequences from public and assembled metagenomic datasets and performed all related analyses. H.W. and Y.Z. performed the assembly of over 20,000 human gut metagenomic samples. L.Y., Y.Q., A.P.C., E.V.K., and L-X.C. drafted the manuscript. K.A. and Y.D. provided suggestions on analyses and commented on the manuscript. All authors finalized and approved the manuscript.

