## Supplementary Figures 1-22 for "A distinct class of conjugative megaplasmids includes potential vehicles for prophage dissemination"

Karthik Anantharaman[3], Haoyu Wang (王浩宇)[4,5], Yuanqiang Zou (邹远强)[4], Yi Duan (段屹)[6,7],

Antonio Pedro Camargo[8,*], Eugene V. Koonin[9,*], LinXing Chen (陈林兴)[1,*]

[1] State Key Laboratory of Advanced Environmental Technology, the Department of Environmental Science and Engineering, University of Science and Technology of China, Hefei, China

[2] Innovative Genomics Institute, University of California, Berkeley, Berkeley, CA 94720, USA

[3] Department of Bacteriology, University of Wisconsin-Madison, Madison, WI, USA

[4] State Key Laboratory of Genome and Multi-omics Technologies, BGI Research, Shenzhen 518083, China

[5] College of Life Sciences, University of Chinese Academy of Sciences, Beijing 100049, China

[6] State Key Laboratory of Immune Response and Immunotherapy, Department of Infectious Diseases, The First Affiliated Hospital of USTC, Center for Advanced Interdisciplinary Science and Biomedicine of IHM, Division of Life Sciences and Medicine, University of Science and Technology of China, Hefei, Anhui 230027, China

[7] Key Laboratory of Anhui Province for Emerging and Reemerging Infectious Diseases, Hefei, Anhui 230027, China

[8] Department of Biochemistry, Institute of Chemistry, University of São Paulo, São Paulo, SP 05508-060, Brazil

[9] Computational Biology Branch, Division of Intramural Research, National Library of Medicine, National Institutes of Health, Bethesda, MD 20894, USA

[*]Corresponding author:

LinXing Chen,

Eugene V. Koonin,

Antonio Pedro Camargo,

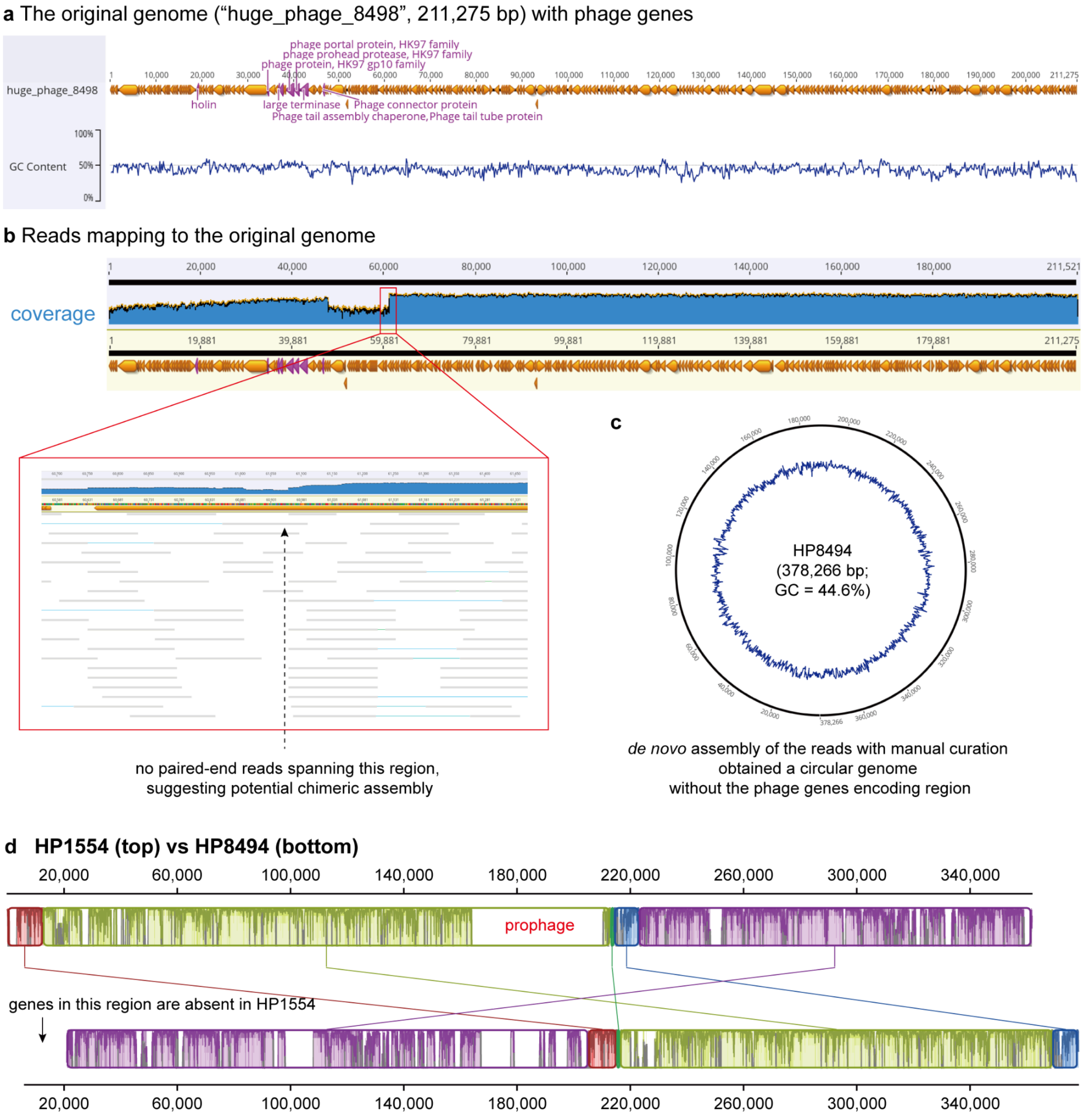


**Supplementary Figure S1 | Genome curation of "huge_phage_8498".** (a) The genome encoded some phage genes. (b) Paired-end reads mapping to the genome suggested potential chimeric assembly issues. (c) The complete and circular genome. (d) The genome-wide comparison of HP8494 (Hodor15) and HP1554 (Hodor10). The genome comparison was performed using Mauve [1] within Geneious Prime [2].

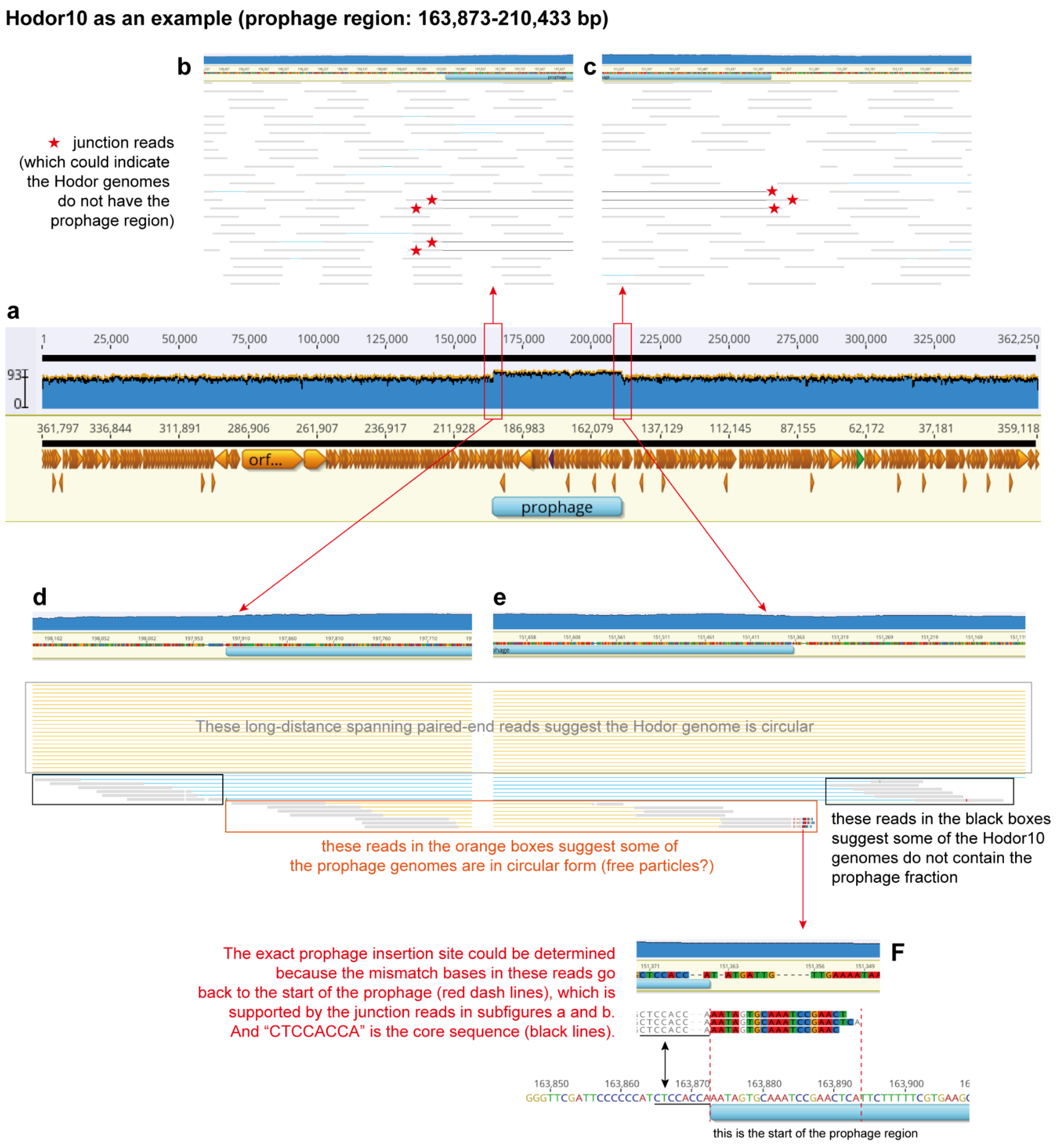


**Supplementary Figure S2 | Some Hodor genomes contain complete prophage genomes.** Hodor10 is used as an example for illustration. (a) Genome-wide view of a representative Hodor genome carrying an integrated prophage. The upper panels show paired-end read mapping across the entire circularized Hodor genome. The coverage track (blue area with black/yellow line) indicates sequencing depth across the replicon. The prophage region is highlighted. The junction reads indicating Hodor genomes without the prophage are enlarged in panels (b) and (c). Long-distance spanning paired-end reads supporting circularization of the Hodor genome are enlarged in panels (d) and (e). (f) Evidence for circularized prophage genomes. Read pairs spanning the prophage termini (orange boxes) suggest that a subset of prophage genomes exists in a circular form, consistent with excision from the Hodor backbone and potential presence in extrachromosomal form and/or phage particles. A high-resolution view of the prophage insertion site is shown. The junction reads (including those highlighted in subpanels a and b) defined the precise prophage integration boundary. Mismatched bases at the read ends map to the start of the prophage region (indicated by red dashed lines), allowing accurate identification of the attachment site. The conserved core sequence "CTCCACCA" (black lines) is present at the insertion junction and represents the recombination core motif.

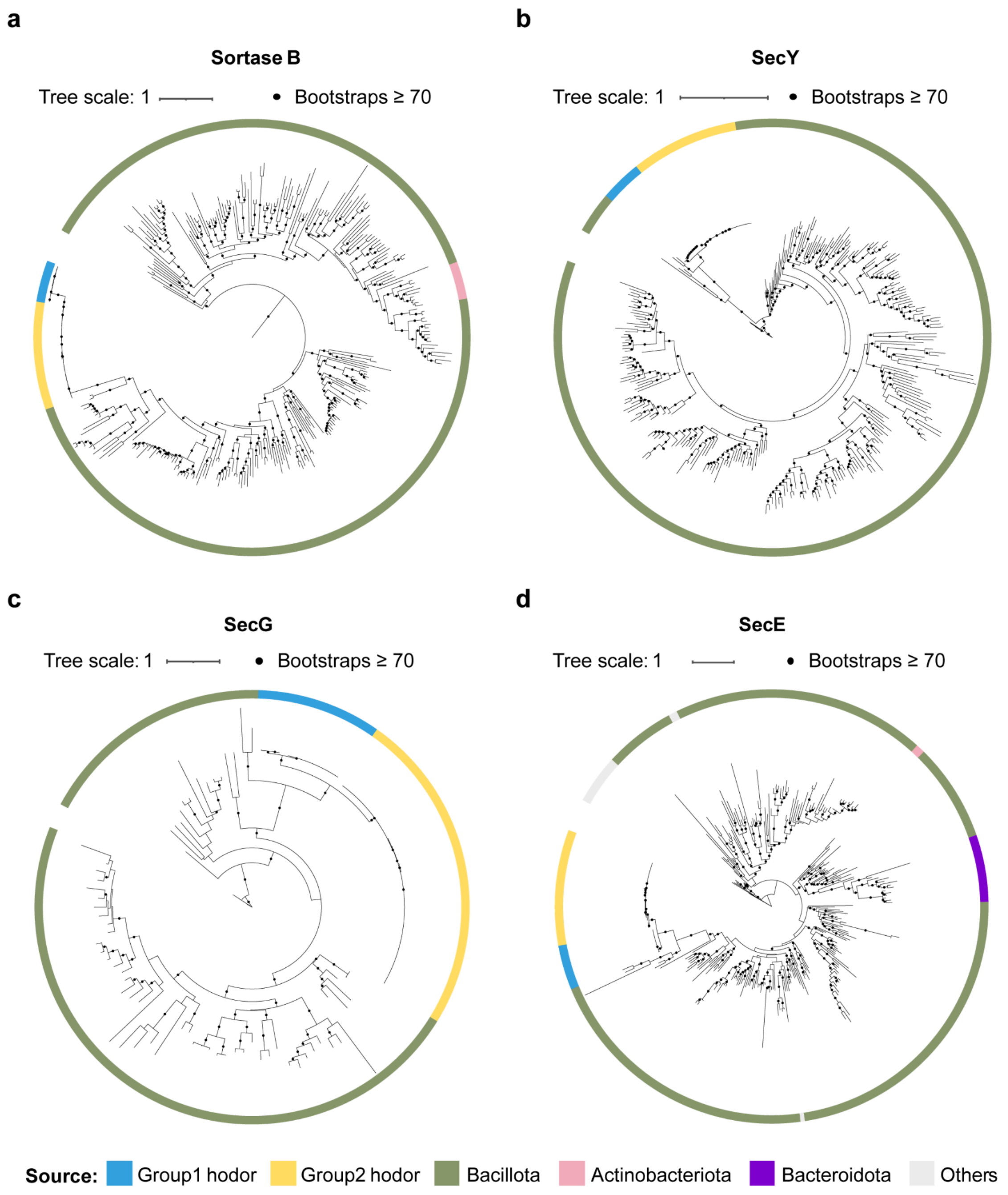


**Supplementary Figure S3 | Hodor genes similar to those in Bacillota.** Maximum-likelihood phylogenetic trees of Hodor-encoding Sortase B (a), SecY (b), SecG (c), and SecE (d), and reference sequences from the HRGMv2_100 protein database. The outer color strip denotes the Hodor group or the taxonomic origin of each sequence.

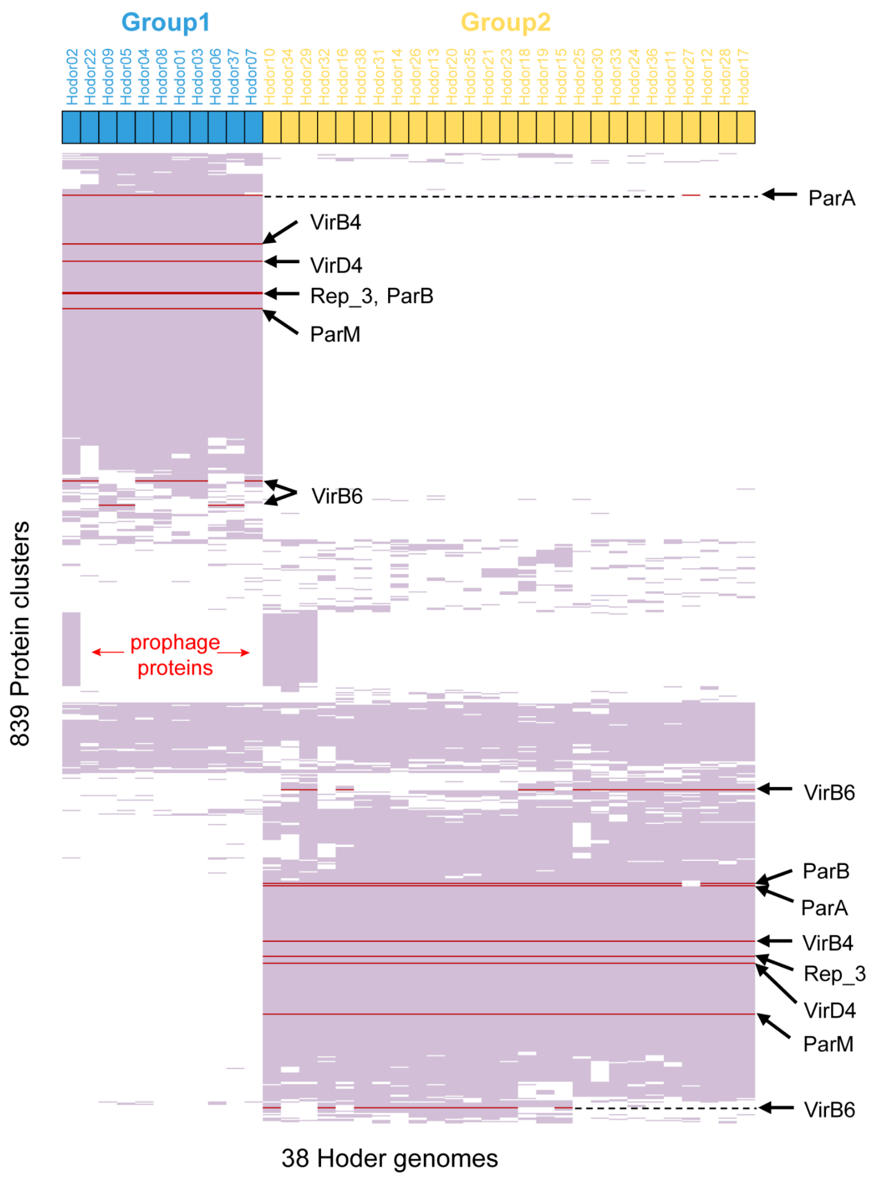


**Supplementary Figure S4 | Protein family clustering analysis of the 38 curated Hodor genomes.** Protein-family distribution across 38 Hodor genomes. Heatmap showing the presence/absence profiles of protein families across 38 Hodor genomes. Rows represent protein families and columns represent genomes. Hodor genomes are colored by the indicated groups (Group1, blue; Group2, yellow). Plasmid marker genes are highlighted in red. Protein families were defined by clustering predicted proteins with mmSeqs2 (--min-seq-id 0.7 -c 0.8 --cov-mode 1 --cluster-mode 2), and only those detected in at least 3 genomes are shown here. The proteins of the prophage identified in one G1 and three G2 Hodor genomes are highlighted. Rows and columns were hierarchically clustered based on Euclidean distance.

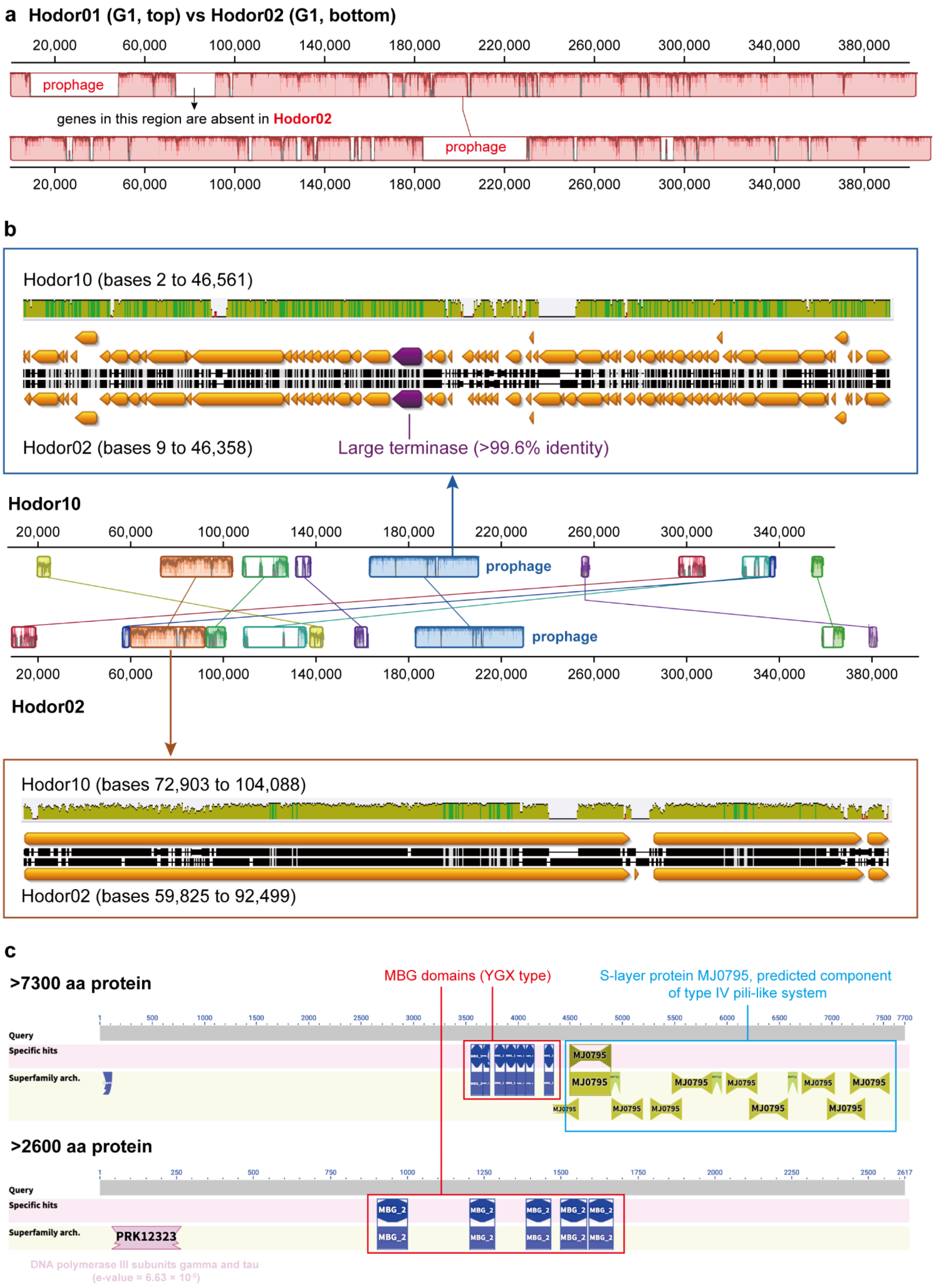


**Supplementary Figure S5 | Detailed information on high similarity regions between Hodor10 and Hodor02.** (a) Hodors from the same group contained different prophages. The genome comparison of Hodor01 and Hodor02 is shown as an example. (b) Hodors with divergent genomes share a similar prophage. The genomes of Hodor10 and Hodor02 belong to different Hodor groups, but the embedded prophage genomes are highly similar. (c) Information on the conserved gene block encoding surface proteins. The genome alignment was performed using Mauve [1] within Geneious Prime Build 2025-03-24 [2].

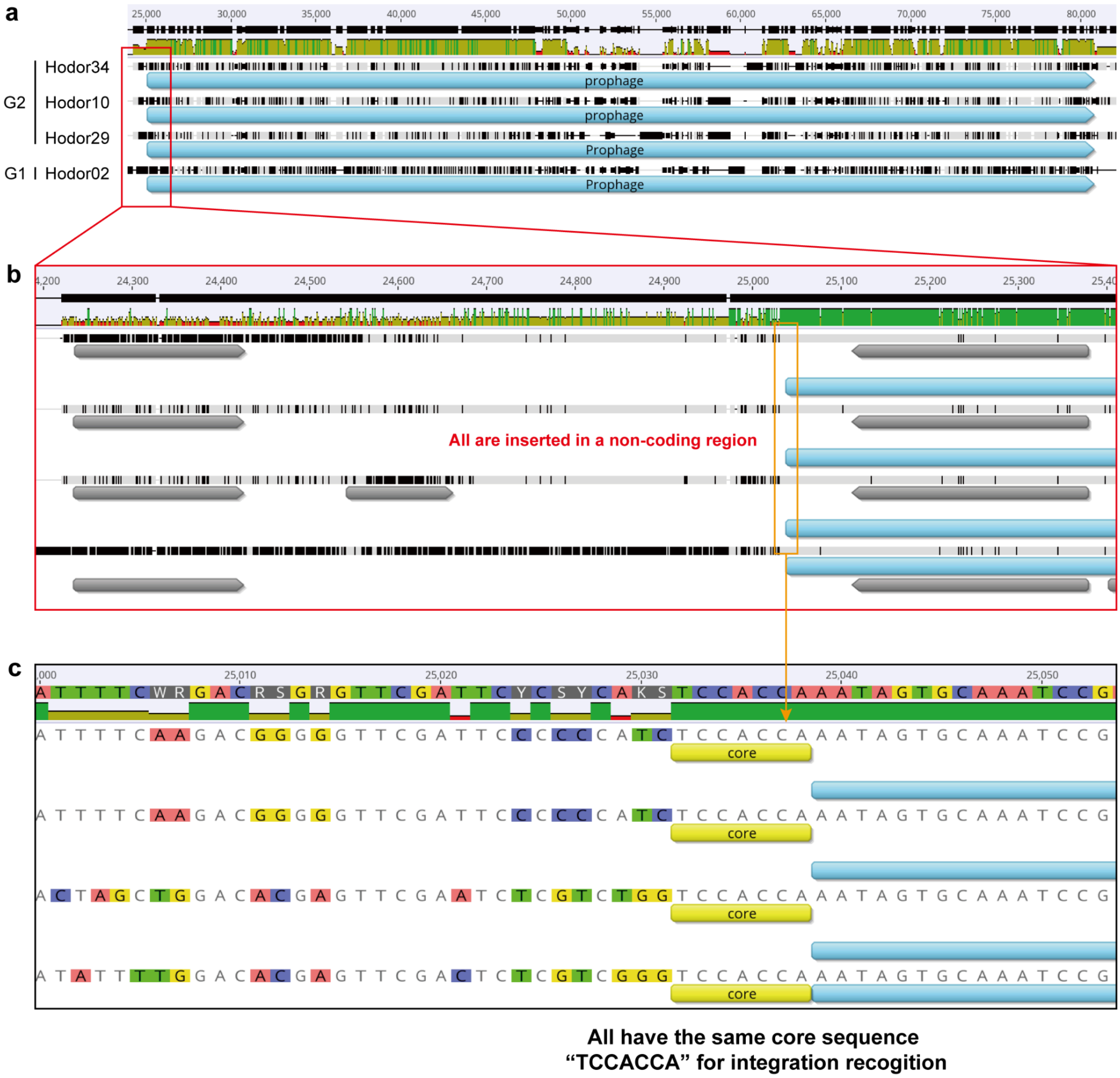


**Supplementary Figure S6 | The conserved integrase location of prophages in Hodor02, 10, 29, and 34.** (a) The prophage regions extracted from the whole Hodor genome alignments. (b) Zoom-in to show that the prophages are all inserted into a non-coding region in the Hodor genomes. (c) Zoom in to show that the core sequences for integration recognition are the same for all prophages.

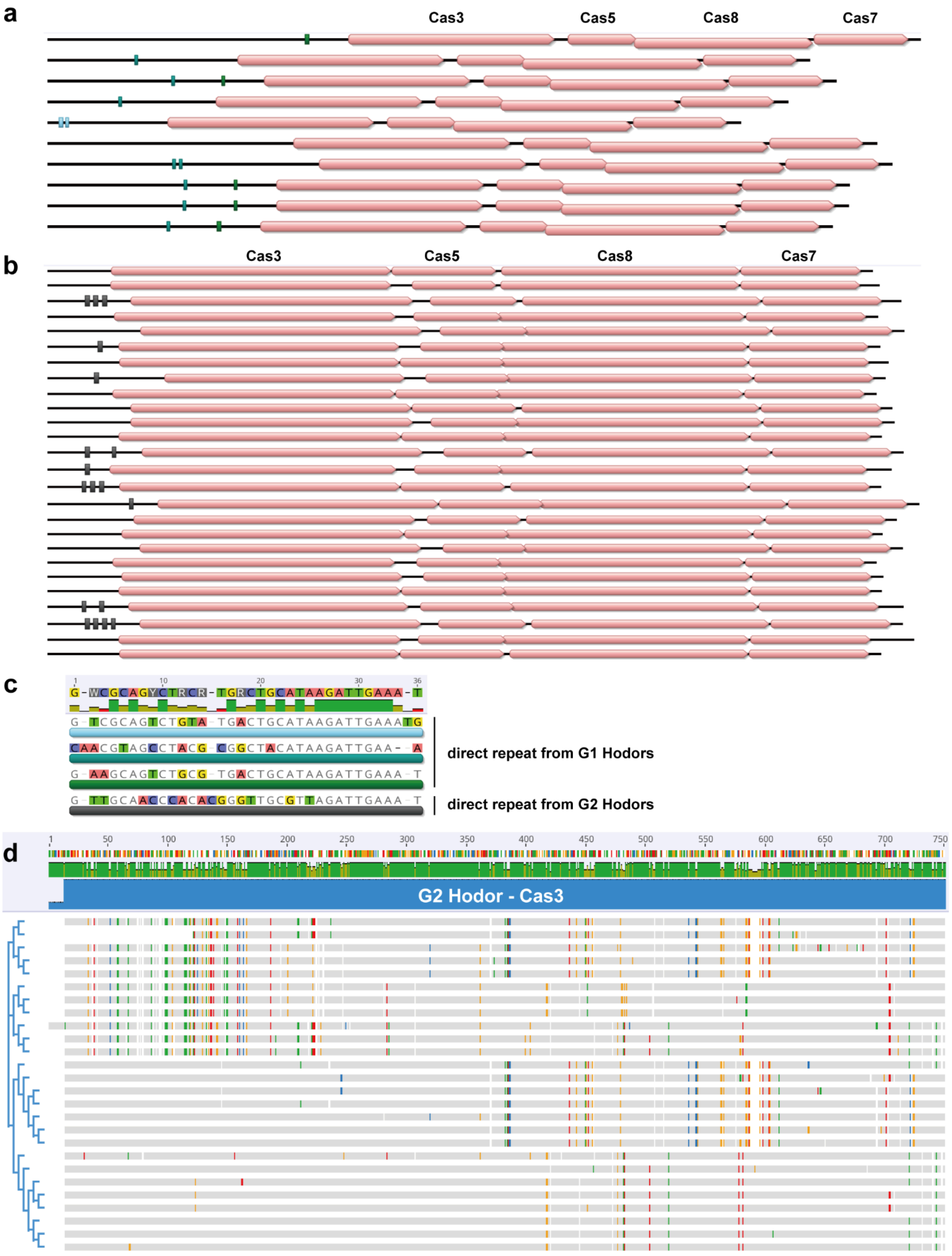


**Supplementary Figure S7 | CRISPR-repeat analyses of Logan retrieved G2 Hodor contigs.** (a) The CRISPR-Cas system in G1 Hodors. (b) The CRISPR-Cas system was in G2 Hodors. In (a) and (b), the squares indicate the predicted direct repeat sequences. (c) Comparison of direct repeat sequences predicted from G1 and G2 Hodors. (d) The phylogeny and alignment of the Cas3 proteins in the CRISPR-Cas systems of G2 Hodors. The Cas3 proteins in G1 CRISPR-Cas systems were almost identical and thus not shown here.

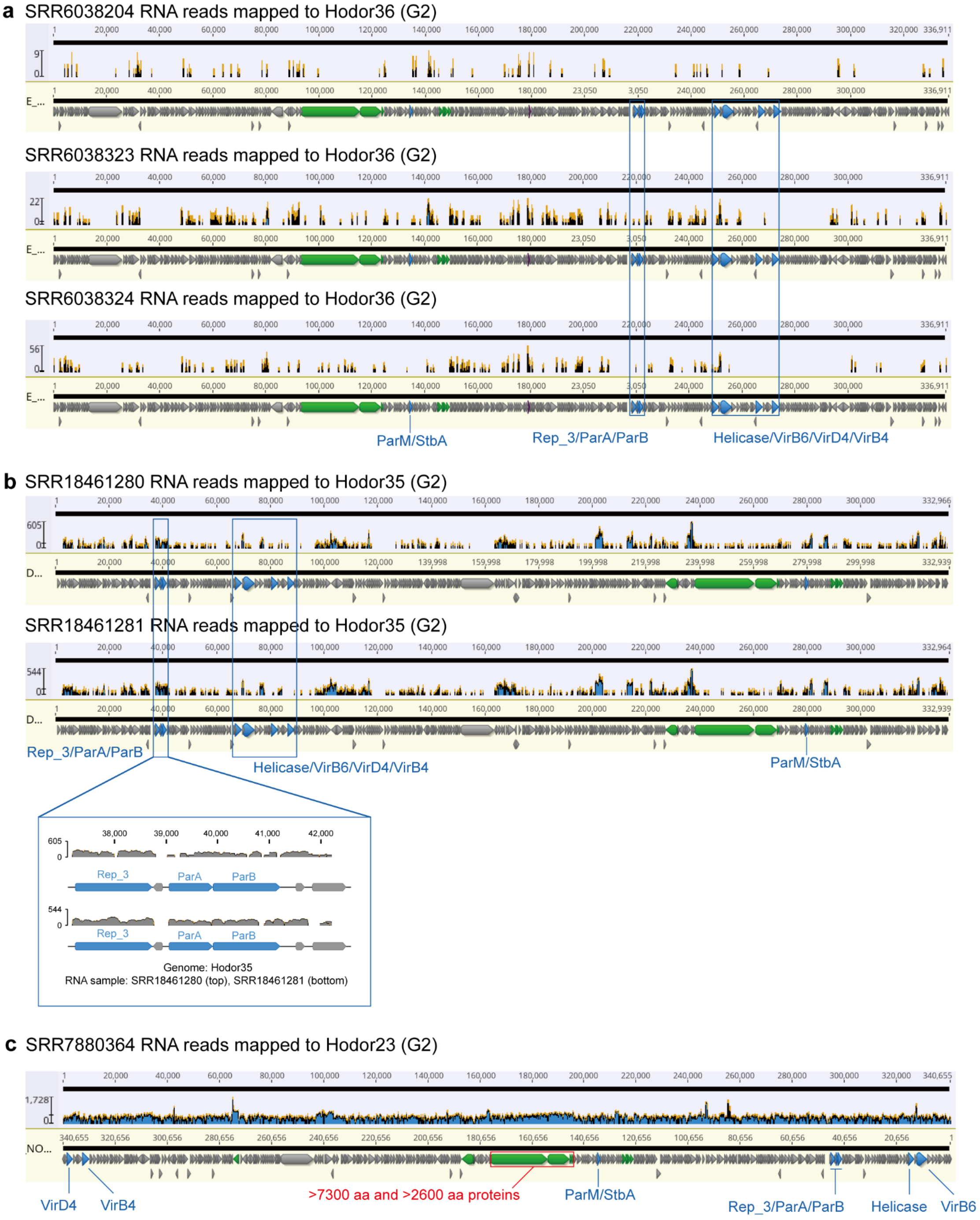


**Supplementary Figure S8 | Transcriptional activities of Hodors.** (a) The RNA reads from three time points were mapped to the Hodor36 genome of G2. (b) The RNA reads mapped to the Hodor35 genome of G2. The Hodor35 genome was reconstructed from the stool sample of a pediatric patient, which is used as an example, along with two corresponding RNA samples, one from a distal colon sample (SRR18461280), the other from a proximal colon sample (SRR18461281) [3]. (c) The RNA reads mapping profile of the Hodor23 genome of G2. The >7300 aa protein and its neighborhood genes are highlighted with a red box. The NCBI SRA id of the RNA dataset is listed above the figure.

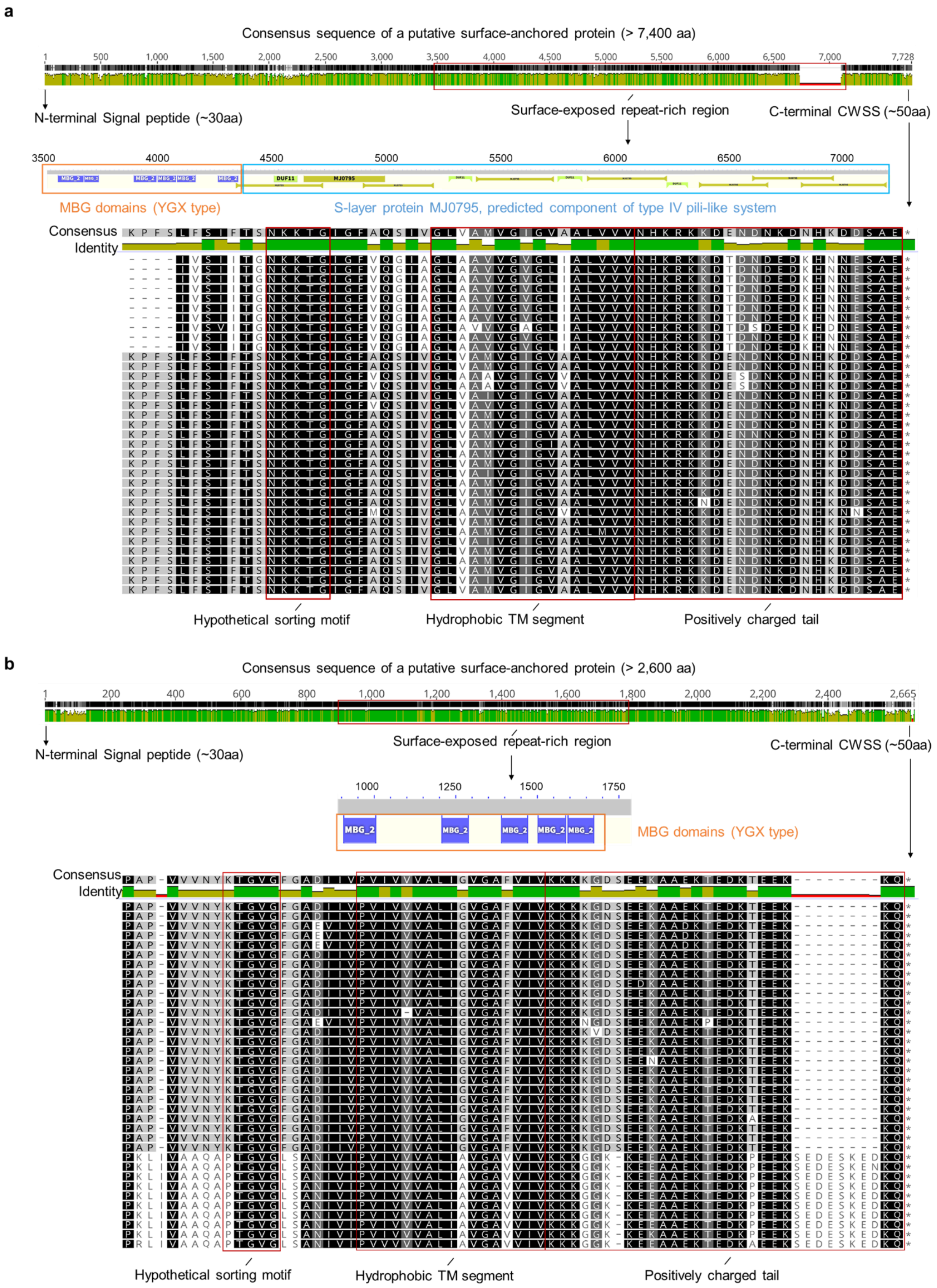


**Supplementary Figure S9 | Architectural features of two putative sortase-anchored surface proteins.** Two large proteins (a and b) with an N-terminal signal peptide, a central surface-exposed repeat-rich region, and a C-terminal cell wall sorting signal (CWSS), consistent with the canonical architecture of sortase-anchored surface proteins. For each protein, multiple-sequence alignments of the C-terminal CWSS region highlight the putative sorting motif, the hydrophobic transmembrane (TM) helix, and the positively charged cytosolic tail. The surface-exposed repeat-rich region was annotated using online BLASTp searches of the consensus sequences.

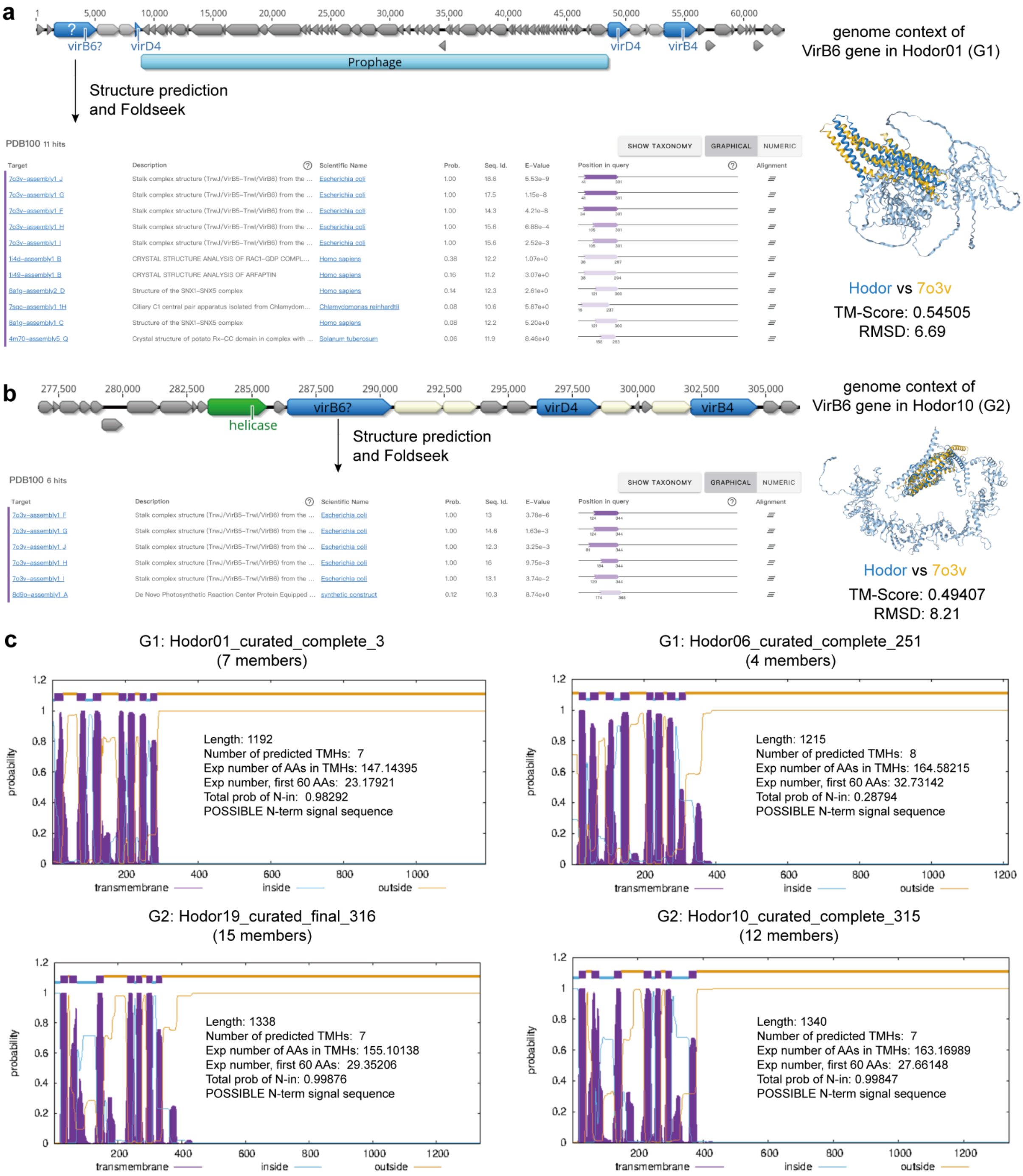


**Supplementary Figure S10 | VirB6 prediction.** The genetic context of the potential VirB6 gene encoded by (a) G1 and (b) G2 Hodors. The foldseek online search results (https://search.foldseek.com/search) and the protein structure alignment of the best hit are shown. Hodor01 from G1 and Hodor10 from G2 are used as examples. Both the predicted G1 and G2 VirB6 were with PDB 7o3v as the structure alignment hit. (c) The transmembrane analyses of the putative VirB6. The details of the transmembrane features are shown. The analyses were performed online with default parameters (https://services.healthtech.dtu.dk/services/TMHMM-2.0/). Note that the VirB6 sequences from the 38 curated Hodor genomes were assigned to four protein families based on ≥70% identity across ≥80% of their length.

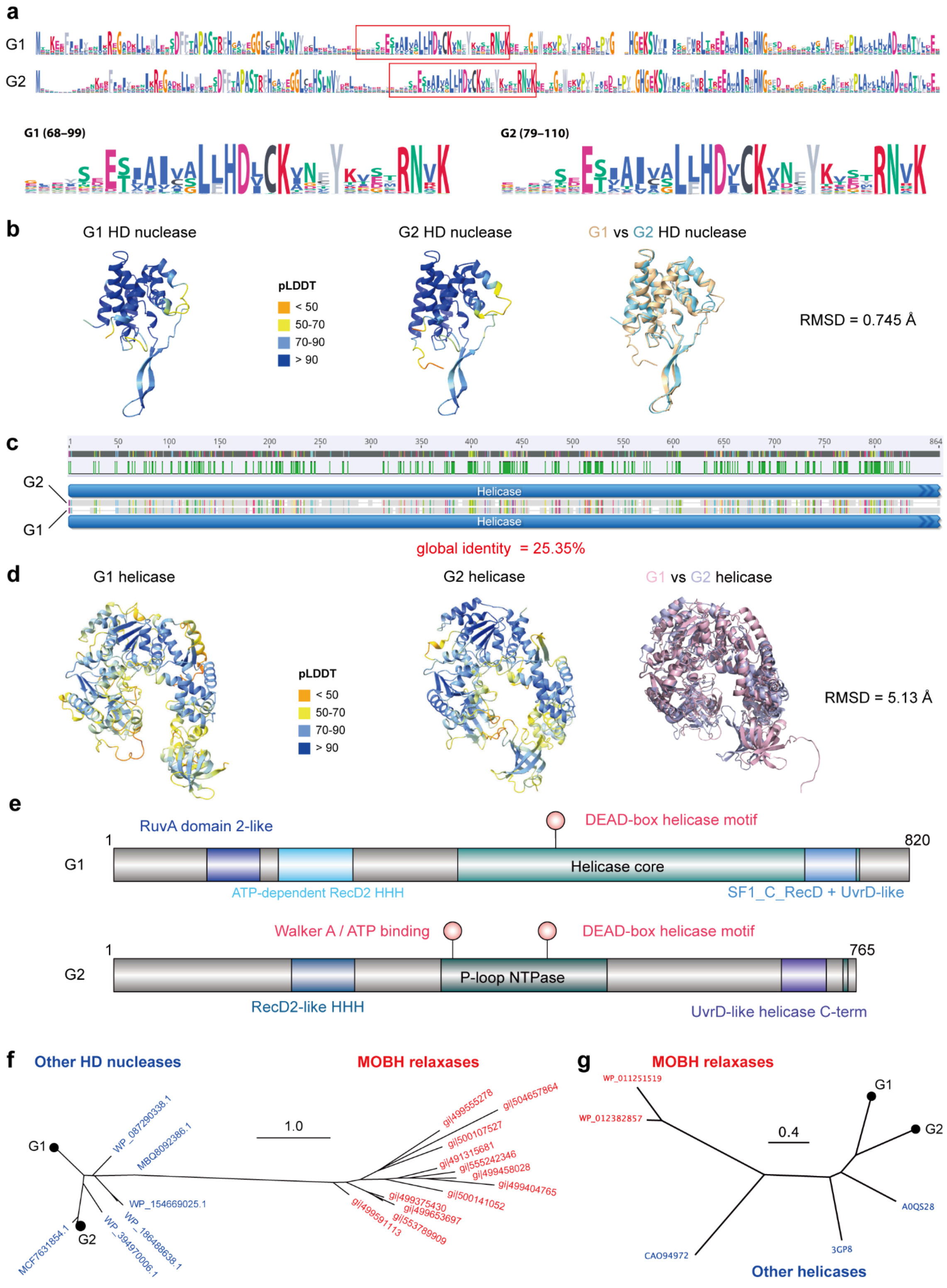


**Supplementary Figure S11 | Analysis of Hodor HD nuclease and helicase proteins.** (a) The sequence logo of the G1 and G2 HD nucleases, the residues in the red boxed are zoomed in. (b) Sequence structure alignment of G1 and G2 HD nucleases. (c) Sequence alignment of G1 and G2 Hodor helicases. (d) Sequence structure alignment of G1 and G2 helicases. (e) Sequence-based functional annotation of G1 and G2 helicase proteins. Conserved helicase-related domains, including the P-loop NTPase core and the UvrD-like helicase C-terminal domain, are shown as colored blocks. Key conserved helicase residues are indicated by symbols, including a Walker A ATP-binding residue, the DEAD-box helicase motif, and a conserved C-terminal ATP-binding residue. (f) The phylogenetic relatedness of Hodor HD nucleases to other HD nucleases (blue) and relaxases of the $MOB_H$ family (red). (g) The phylogenetic relatedness of Hodor helicases to other helicases (blue) and relaxases of the $MOB_H$ family (red).

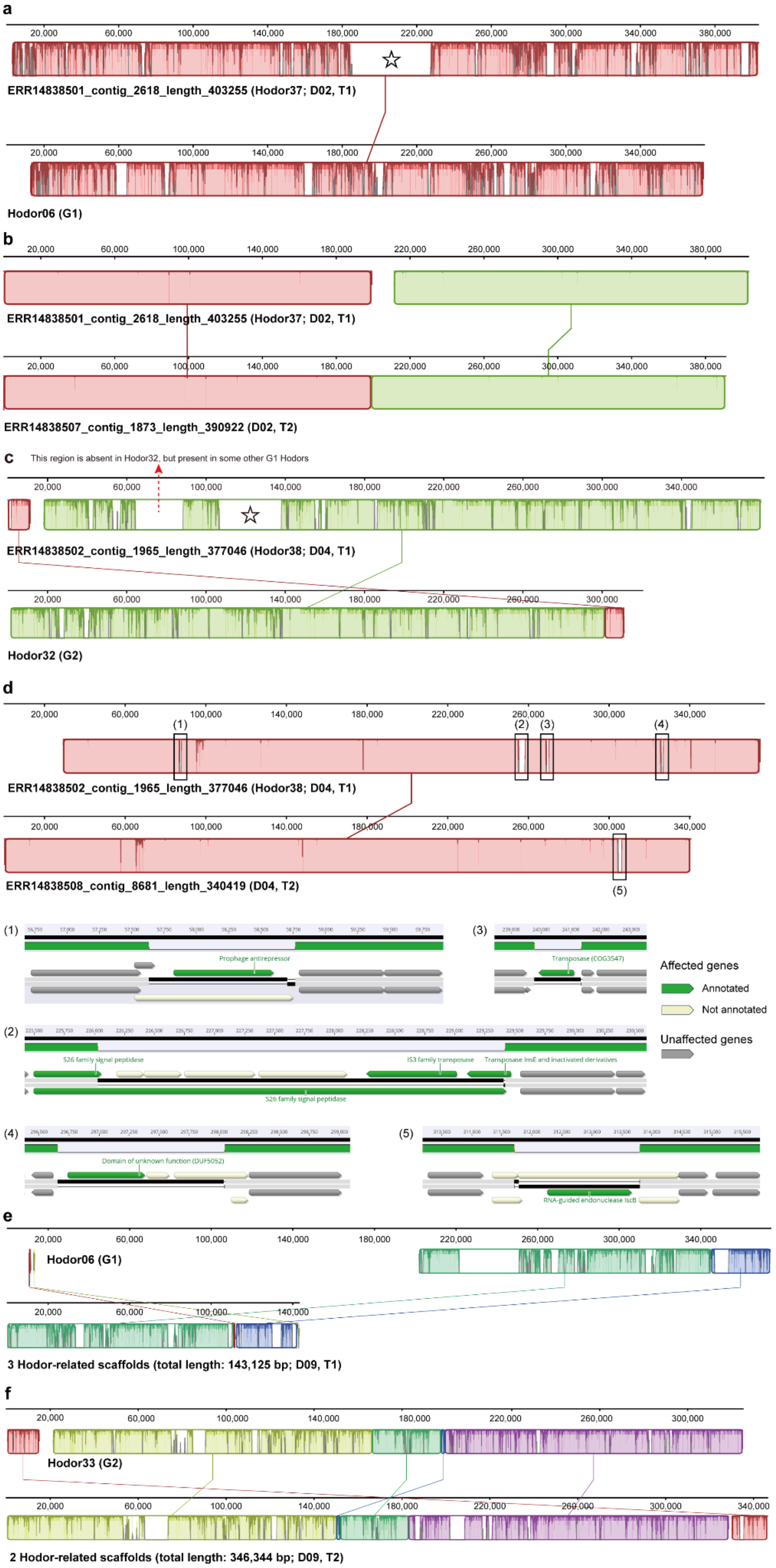
a
ERR14838501_contig_2618_length_403255 (Hodor37; D02, T1)
Hodor06 (G1)
b
ERR14838501_contig_2618_length_403255 (Hodor37; D02, T1)
ERR14838507_contig_1873_length_390922 (D02, T2)
c
This region is absent in Hodor32, but present in some other G1 Hodors
ERR14838502_contig_1965_length_377046 (Hodor38; D04, T1)
Hodor32 (G2)
d
ERR14838502_contig_1965_length_377046 (Hodor38; D04, T1)
ERR14838508_contig_8681_length_340419 (D04, T2)
Prophage antirepressor
Transposase (COG3547)
S26 family signal peptidase
IS3 family transposase
Transposase InsE and inactivated derivatives
Domain of unknown function (DUF5052)
RNA-guided endonuclease IscB
Affected genes
Annotated
Not annotated
Unaffected genes
e
Hodor06 (G1)
3 Hodor-related scaffolds (total length: 143,125 bp; D09, T1)
f
Hodor33 (G2)
2 Hodor-related scaffolds (total length: 346,344 bp; D09, T2)

**Supplementary Figure S12 | The detection and comparison of Hodor genomes in metagenomic assemblies of the long-read datasets.** (a) The Hodor genome detected in the sample of individual D02 collected at the first time (T1) is most similar to the curated genome of Hodor06 from G1. (b) The Hodors in D02 collected from both time points (T1 and T2) are almost identical. (c) The Hodor genome detected in the sample of D04 from T1 is most similar to the curated genome of Hodor32 from G2. (d) The Hodors in D04 collected from T1 and T2 have highly similar genomes, with several gene gain and lose events. In detail, the Hodor collected at T2 lacks genes for (1) a prophage antirepressor, (2) a transposase, (3) two transposases and four hypothetical proteins, (4) a DUF5052 protein and two hypothetical proteins, while the Hodor collected at T1 lacks (5) an RNA-guided endonuclease IscB gene. (e) The Hodor in D09 from T1 is most similar to the curated genome of Hodor06 from G1. (f) The Hodor in D09 from T2 is most similar to the curated genome of Hodor33 from G2. The long-read datasets were reported by Wirbel et al. (Wirbel et al., 2026). The genome alignment was performed using Mauve [1] within Geneious Prime Build 2025-03-24 [2]. In (a) and (c), the region marked by a pentagon indicates additional complete conjugative apparatus acquired in these genomes.

a

Hodor37 (G1; 403,253 bp in length; D02, T1)

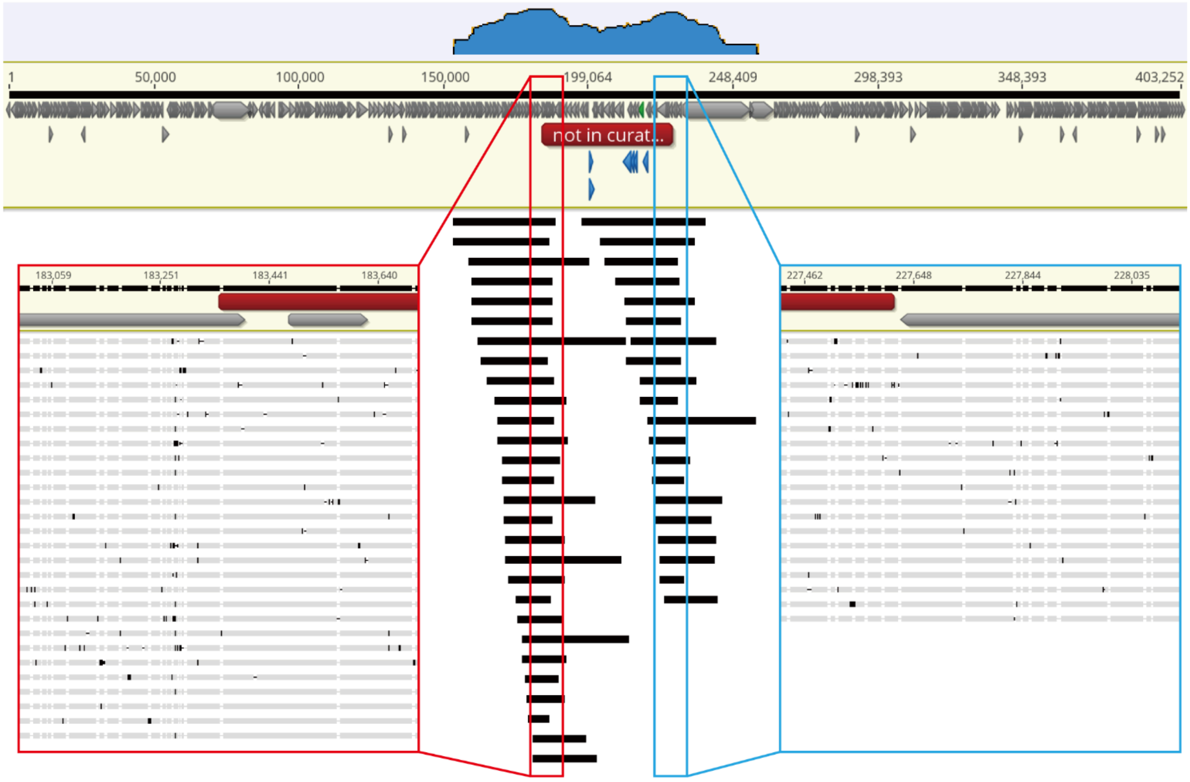


b

Hodor38 (G2; 379,934 bp in length; D04, T1)

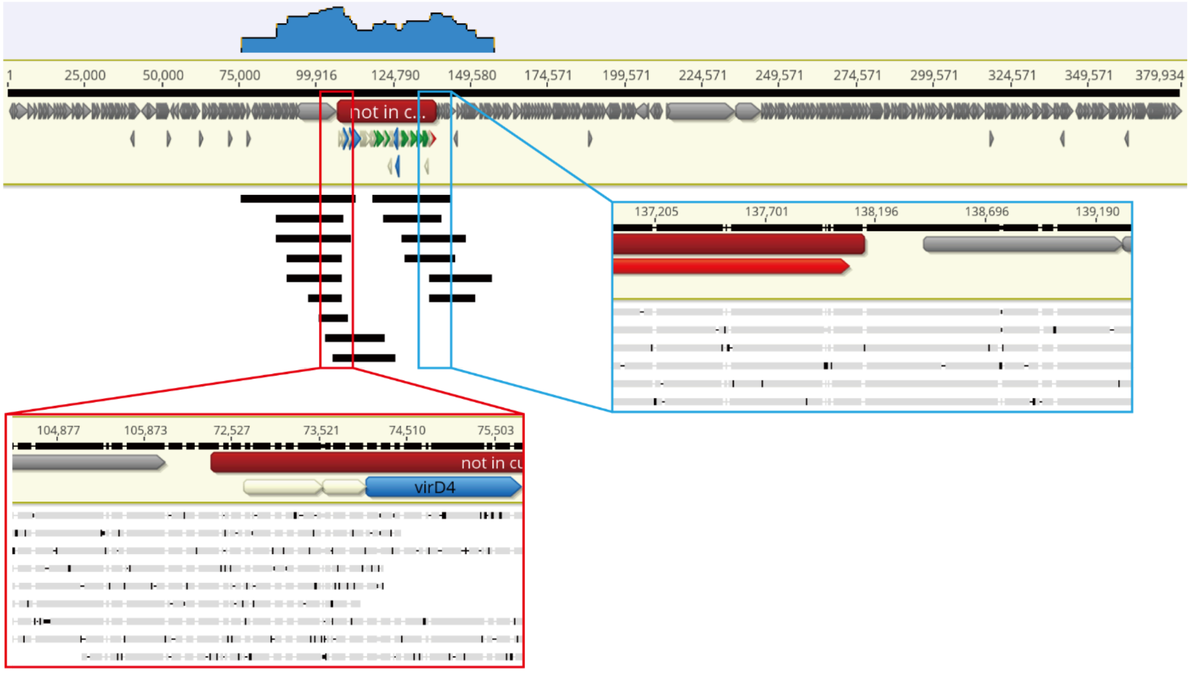


**Supplementary Figure S13 | The detection of complete conjugative systems in two additional curated Hodor genomes is not due to assembly errors.** We retrieved the long reads by BLASTn and mapped them to the region spanning the potential insertion location, and all mapped reads support the assembly. Note that the frequent SNPs are normal, which are due to the indels in long read sequencing. (a) The Hodor genome (Hodor37, which belongs to G1) assembled from the individual D02 at the first sampling point (T1). (b) The Hodor genome (Hodor38, belongs to G2) assembled from the individual D04 at the first sampling point (T1).

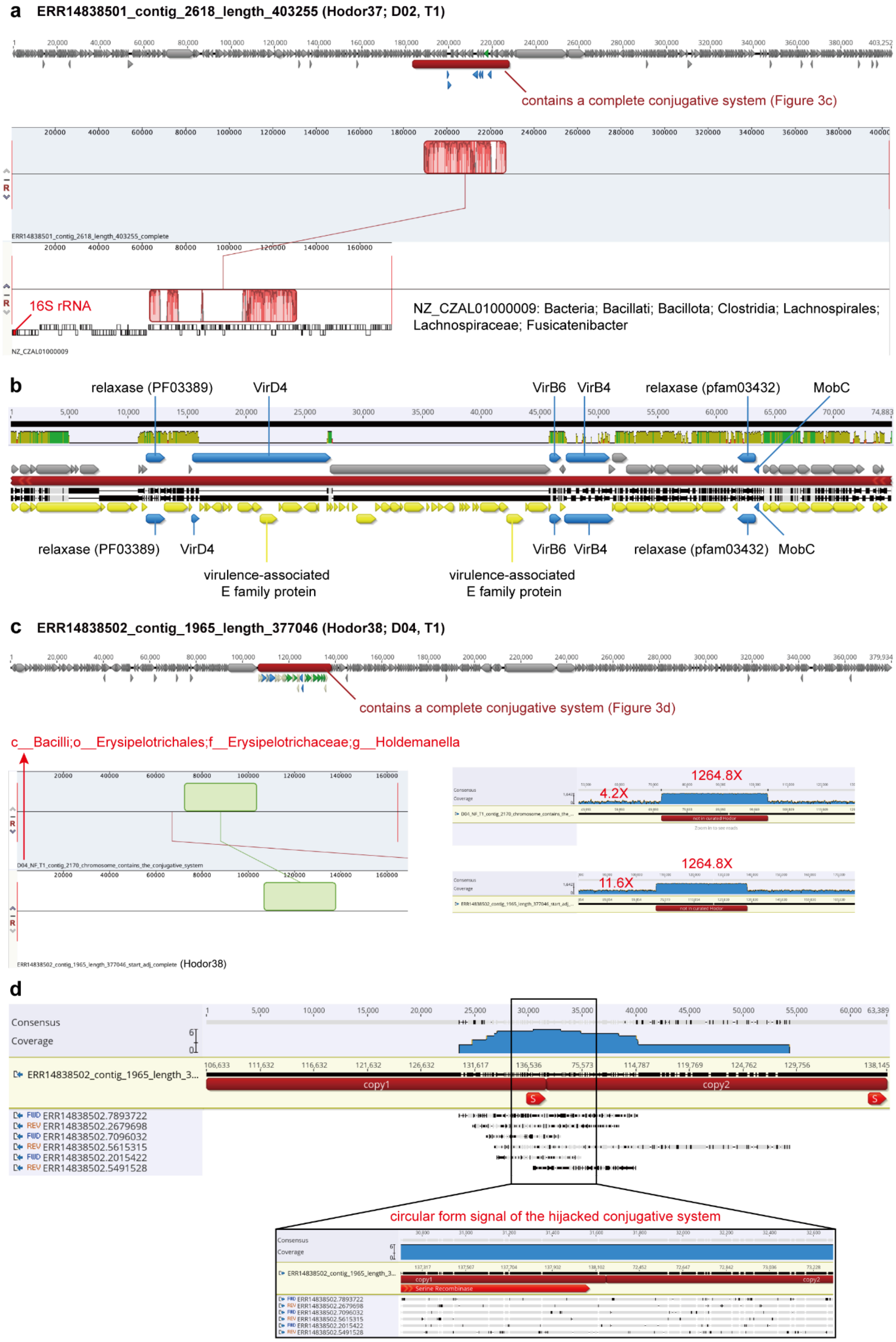
a
ERR14838501_contig_2618_length_403255 (Hodor37; D02, T1)
contains a complete conjugative system (Figure 3c)
ERR14838501_contig_2618_length_403255_complete
16S rRNA
NZ_CZAL01000009: Bacteria; Bacillati; Bacillota; Clostridia; Lachnospirales; Lachnospiraceae; Fusicatenibacter
NZ_CZAL01000009
b
relaxase (PF03389)
VirD4
VirB6
VirB4
relaxase (pfam03432)
MobC
virulence-associated E family protein
virulence-associated E family protein
c
ERR14838502_contig_1965_length_377046 (Hodor38; D04, T1)
contains a complete conjugative system (Figure 3d)
c__Bacilli;o__Erysipelotrichales;f__Erysipelotrichaceae;g__Holdemanella
D04_NF_T1_contig_2170_chromosome_contains_the_conjugative_system
ERR14838502_contig_1965_length_377046_start_adj_complete (Hodor38)
1264.8X
4.2X
1264.8X
11.6X
d
Consensus
Coverage
ERR14838502_contig_1965_length_3...
copy1
copy2
FWD ERR14838502.7893722
REV ERR14838502.2679698
FWD ERR14838502.7096032
REV ERR14838502.5615315
FWD ERR14838502.2015422
REV ERR14838502.5491528
circular form signal of the hijacked conjugative system
Serine Recombinase

**Supplementary Figure 14 | Chromosomal ICE-like conjugative systems captured by Hodor37 and Hodor38.** (a) The additional conjugative module present in Hodor37 exhibits high sequence similarity to an ICE-like region within the NCBI contig NZ_CZAL01000009, assigned to the genus Fusicatenibacter (Bacillota). The contig harbors a full-length 16S rRNA gene, showing 99.85% identity to Fusicatenibacter saccharivorans strain HT03-11 (NR_114326.1), supporting its taxonomic assignment. (b) Gene synteny comparison between the Hodor37 conjugative region and the ICE-like region in NZ_CZAL01000009. The conserved plasmid mobility genes are highlighted, including relaxase (PF03432) and MobC homologs, which share ≥95.7% and ≥97.2% amino acid identity, respectively. Two genes encoding virulence-associated E family proteins from NZ_CZAL01000009 are indicated. (c) The conjugative module in Hodor38 is identical to a region from a 164,710-bp scaffold assembled from long-read sequencing of the same sample, taxonomically assigned to the genus Holdemanella (Bacillota). Read mapping indicates markedly elevated sequencing depth across the conjugative region (~1264.8×) relative to both the host scaffold (~4.2×) and the Hodor scaffold (~11.6×), suggesting that this module exists in additional genetic contexts beyond either element. (d) Evidence for a circular intermediate of the conjugative module. Two tandem copies of the region were concatenated to detect reads spanning the junction. Six long sequencing reads traversed the artificial boundary, consistent with the presence of an extrachromosomal circular form of this genetic fraction in the sample.

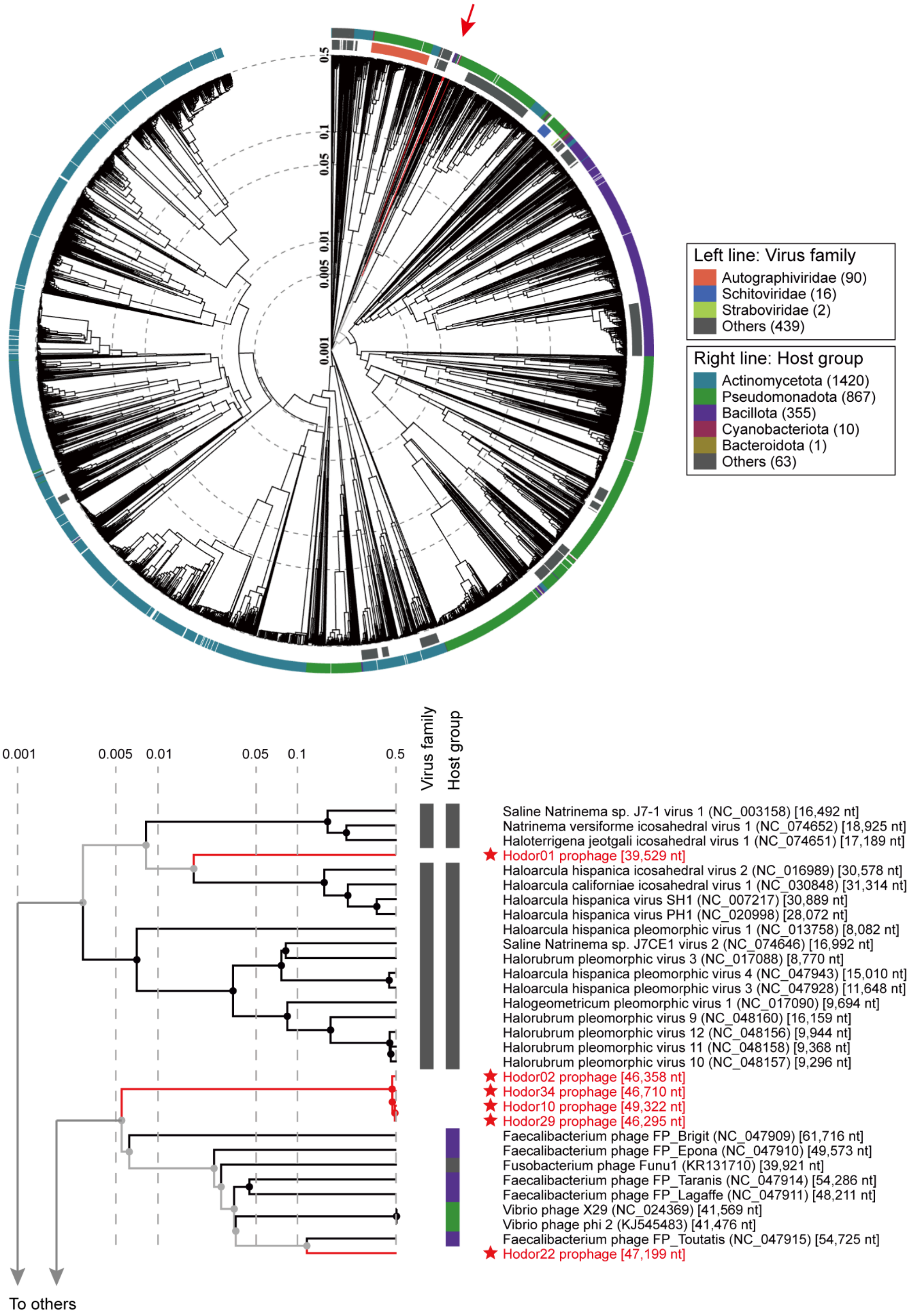


**Supplementary Figure S15 | The taxonomy and potential hosts of Hodor-associated prophages.** The Hodor-associated prophages were uploaded to viptree (https://www.genome.jp/viptree/) [4] for analyses based on the protein families shared with reference dsDNA phages in the viptree database. The upper panel shows the position (red arrow) of the Hodor-associated prophages (highlighted in red and stars), with the genome length (nt) shown in brackets. The bottom panel indicates the closely related phages and their taxonomy and bacterial hosts.

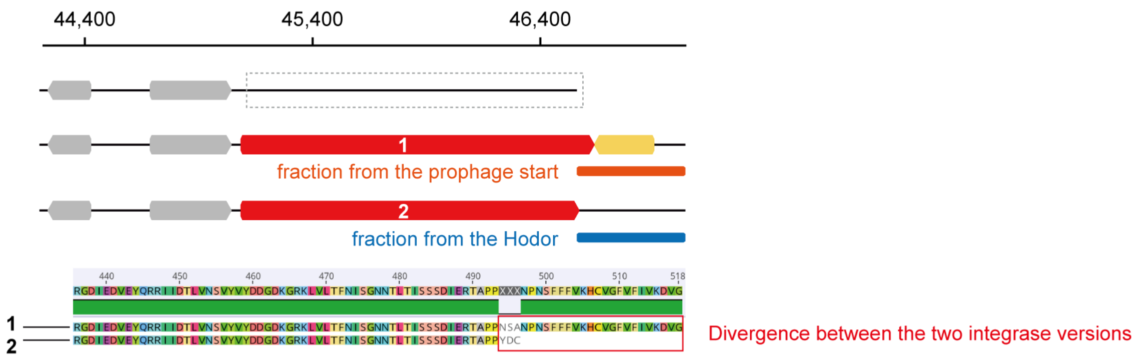


**Supplementary Figure 16 | The integrase genes in prophage01.** The gene could not be predicted in the linear prophage genome (the top illustration), unless a small fraction from the start of the prophage (the middle illustration) or the Hodor genome (the bottom illustration) is added to the end of the linear prophage genome. The red box indicates the divergence between the two versions of the integrase genes.

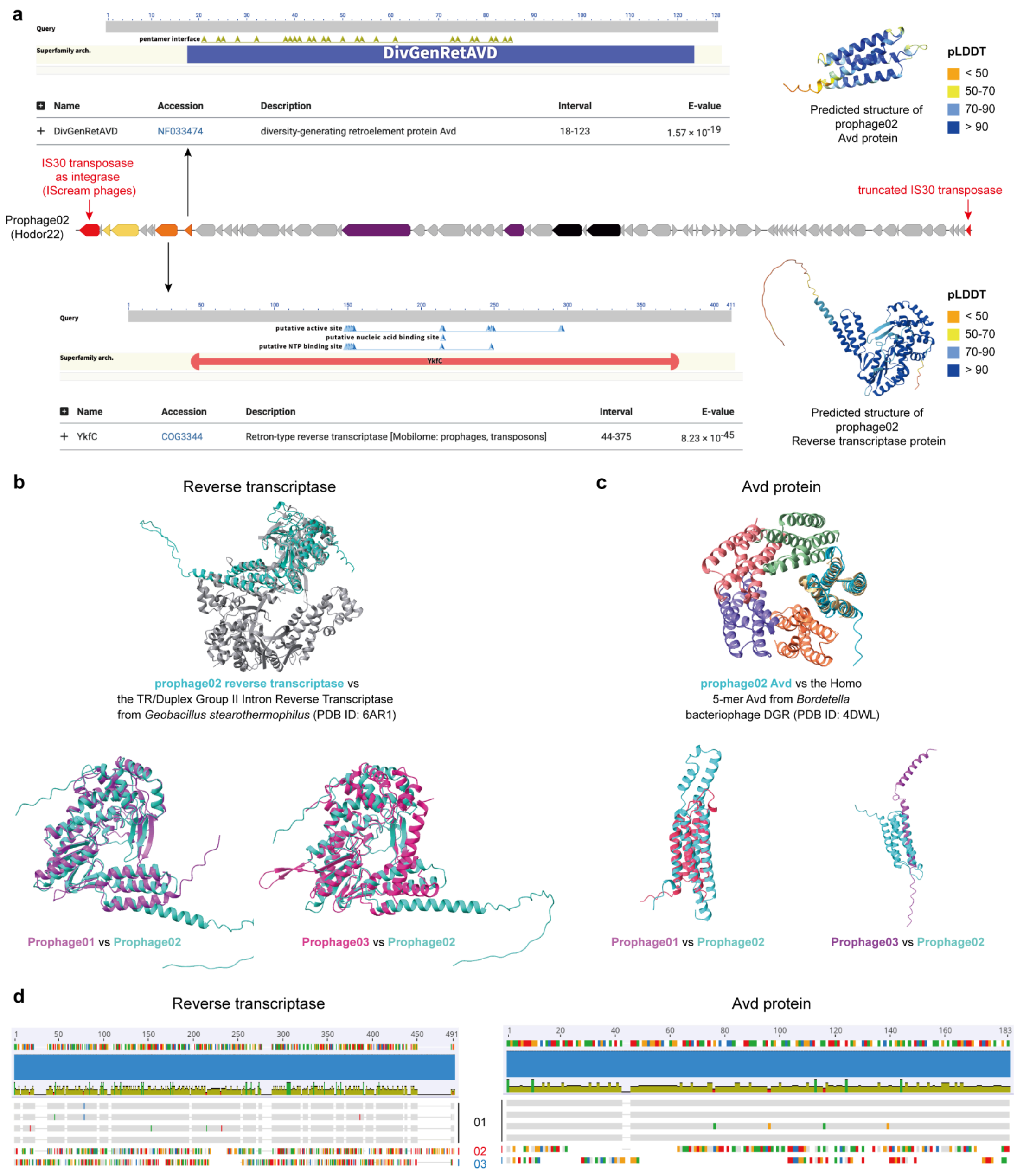


**Supplementary Figure S17 | Reverse transcriptase and Avd proteins identified in Hodor-associated prophages.** (a) The identification of the reverse transcriptase and the Avd protein in prophage02. The protein structures of the predicted genes are shown to the right. (b) The reverse transcriptase annotation in prophage02 was confirmed by structure alignment (vs PDB 6AR1). A similar reverse transcriptase was identified in prophage01 and prophage03. (c) The Avd annotation in prophage02 was confirmed by structure alignment (vs PDB 4DWL). A similar Avd protein was identified in prophage01 and prophage03. (d) Comparison of the reverse transcriptase and the Avd proteins in all three prophage groups.

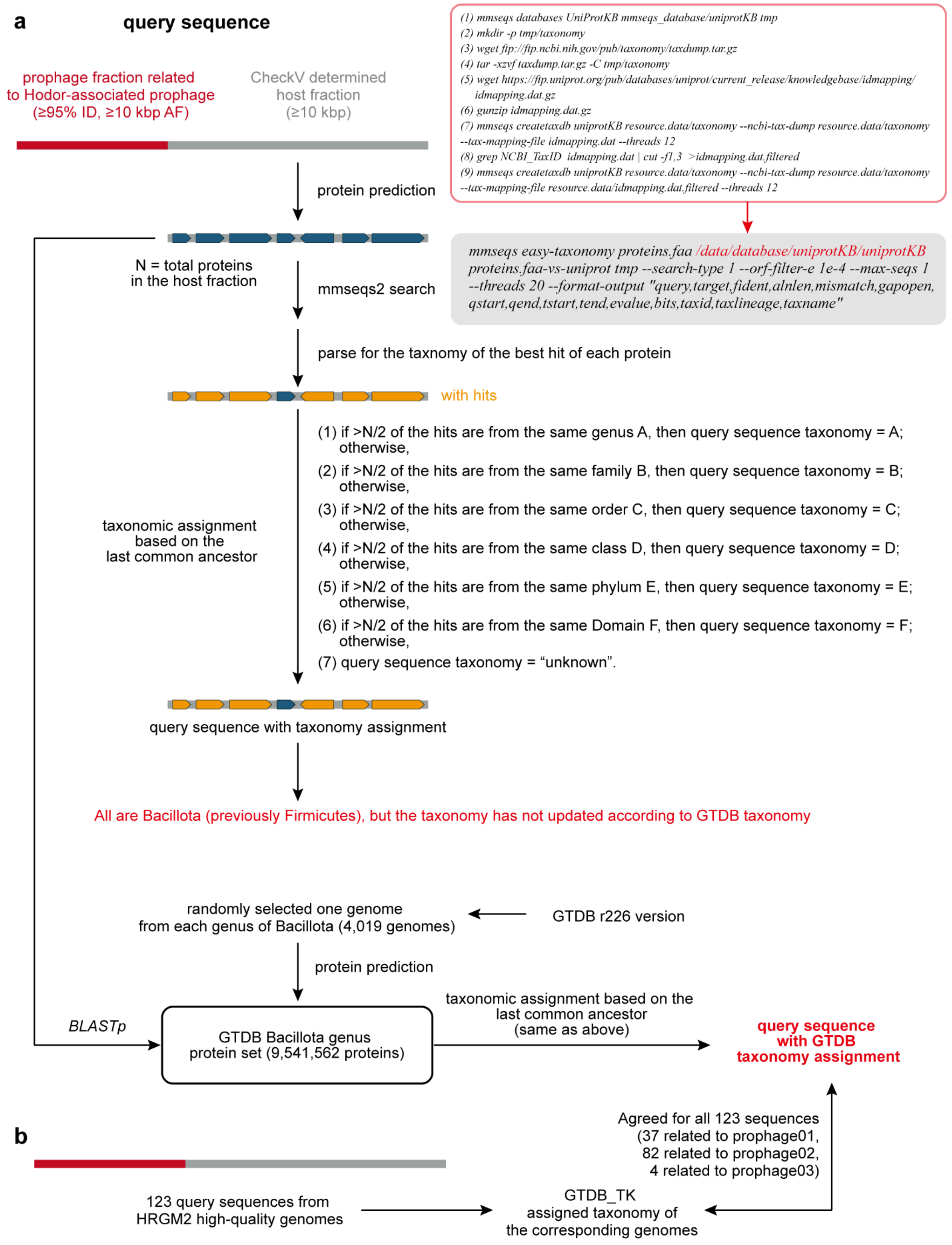


**Supplementary Figure S18 | Taxonomic assignment of host-derived regions flanking Hodor-associated prophages.** (a) Prophage-related regions (≥95% nucleotide identity across ≥10 kb alignment fraction) were first delineated from host genomic fragments identified by CheckV (host fraction ≥10 kb). The protein-coding genes were predicted from the host-associated regions and searched against UniProtKB using mmseqs easy-taxonomy (search-type 1, E-value ≤1e-4, best hit retained). Only proteins with significant matches were retained for downstream taxonomic inference. The resulting hits were assigned taxonomy based on UniProt lineage information (based on NCBI taxonomy) and subsequently reconciled with GTDB classification. Across all examined fragments, taxonomic affiliations consistently mapped to members of Bacillota (formerly Firmicutes), although UniProt annotations have not been fully updated to reflect the current GTDB taxonomy. (b) The taxonomic assignment approaches were approved by the Hodor-associated prophage containing scaffolds from the HRGM2 genomes.

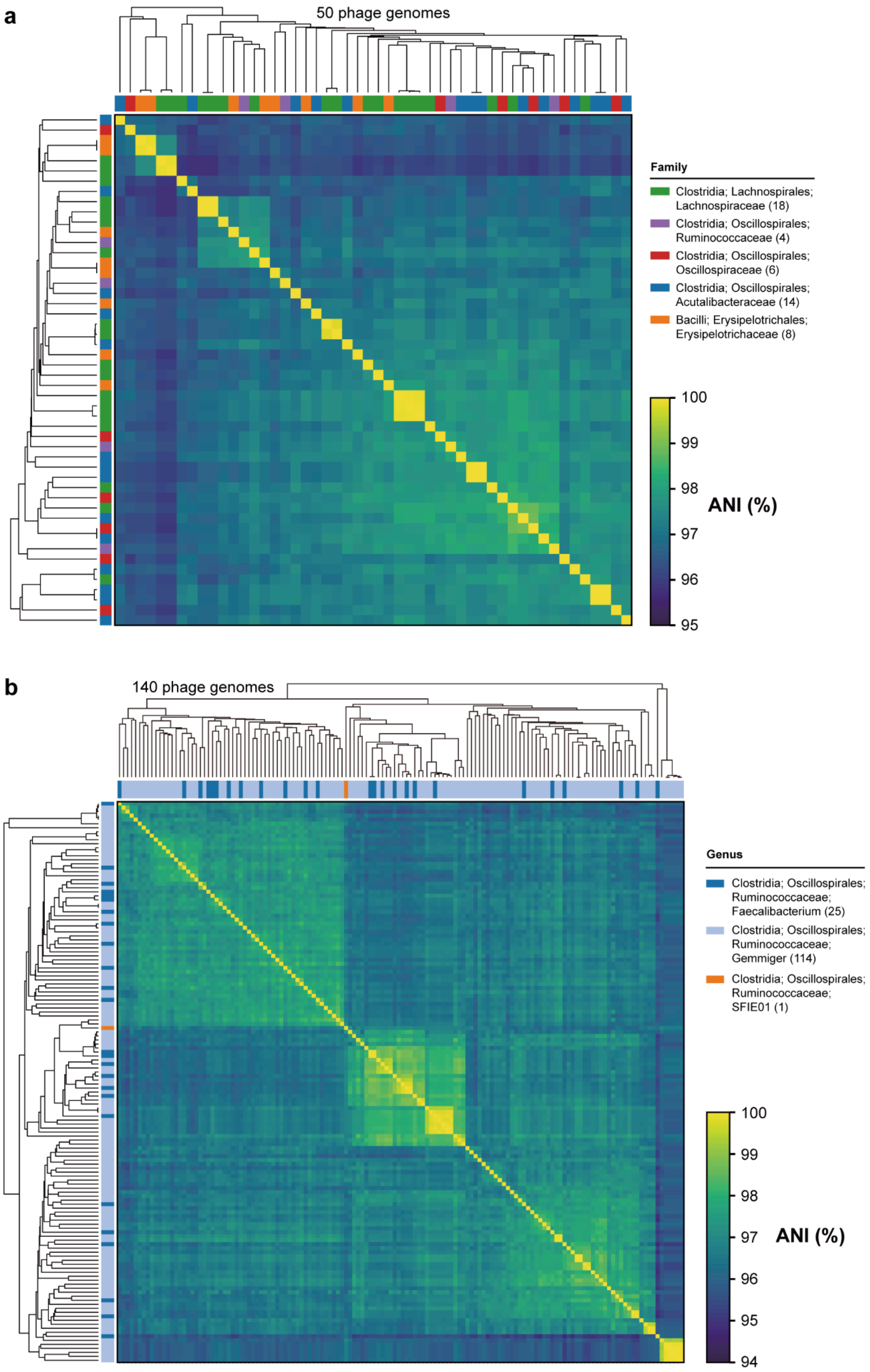


**Supplementary Figure S19 | Average nucleotide identity (ANI) of within-cluster phage sequences.** (a) The Hodor-associated prophage01-related cluster (cluster01a in Figure 5d of the main text). All the phage sequences with host information to the family level and with a minimum length of 40 kbp are included (50 genomes). (b) The Hodor-associated prophage02-related cluster (cluster02 in Figure 5d of the main text). All the phage sequences with host information to the genus level within the family of Ruminococcaceae and with a minimum length of 40 kbp are included (140 genomes). For each taxon, the number of phage genomes is shown in brackets. The ANI values were extracted from the phage sequence clustering analysis on prophage01 and prophage02, respectively, which was based on an all-vs-all BLASTn search (see Methods in the main text).

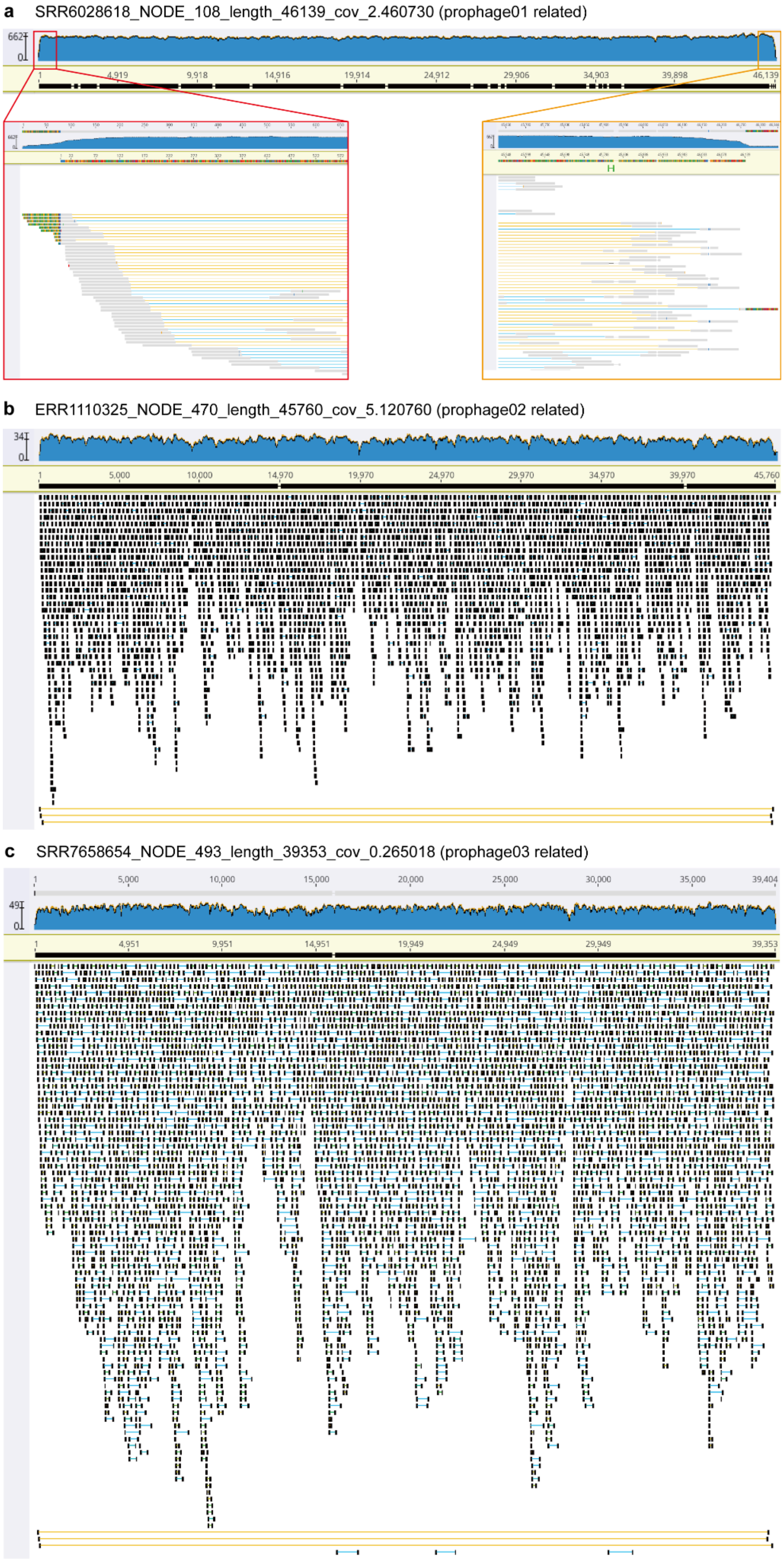


**Supplementary Figure 20 | Reads mapping evidence of Hodor-associated prophages in lytic form.** The paired-end metagenomic reads mapped to the genomes of (a) prophage01, (b) prophage02, and (c) prophage03. The paired-end reads linked by a long yellow line indicate the genome is in circular form.

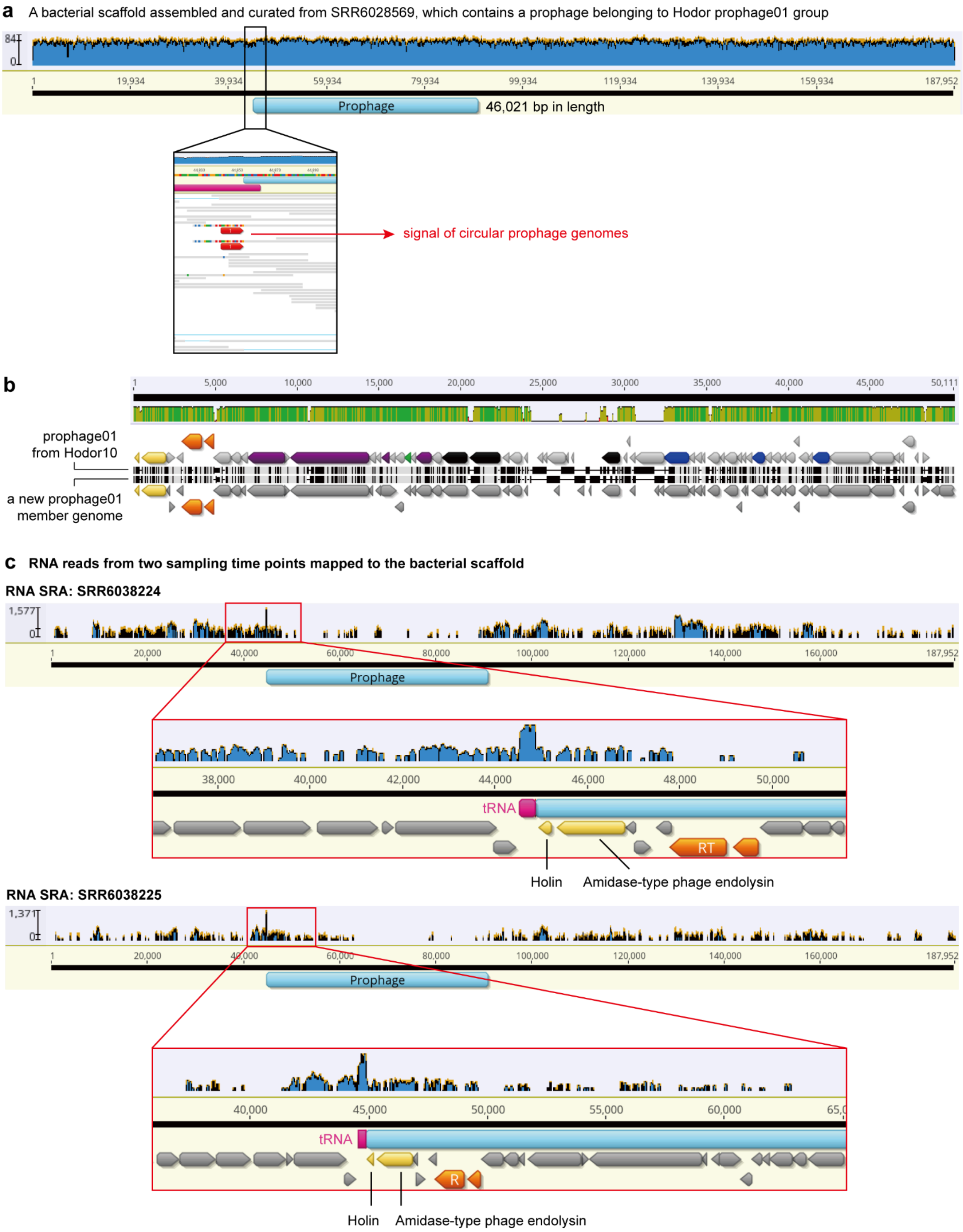


**Supplementary Figure S21 | Metatranscriptomic analysis to show the transcription of lysis-related genes.** (a) The metagenomic reads mapping profiles of the prophage01 member genome. The zoom-in shows the signal of circular prophage genomes (only a small subset, though) in the sample. (b) The genome alignment of the prophage genome to the prophage genome in Hodor10 (G2). (c) The metatranscriptomic reads mapping profiles of the prophage01 member genome. Two metatranscriptomic SRA samples are shown. The zoom-in shows the transcriptional details, with the lysis-related gene names shown. The genome alignment was performed using Mauve [1] within Geneious Prime Build 2025-03-24 [2].

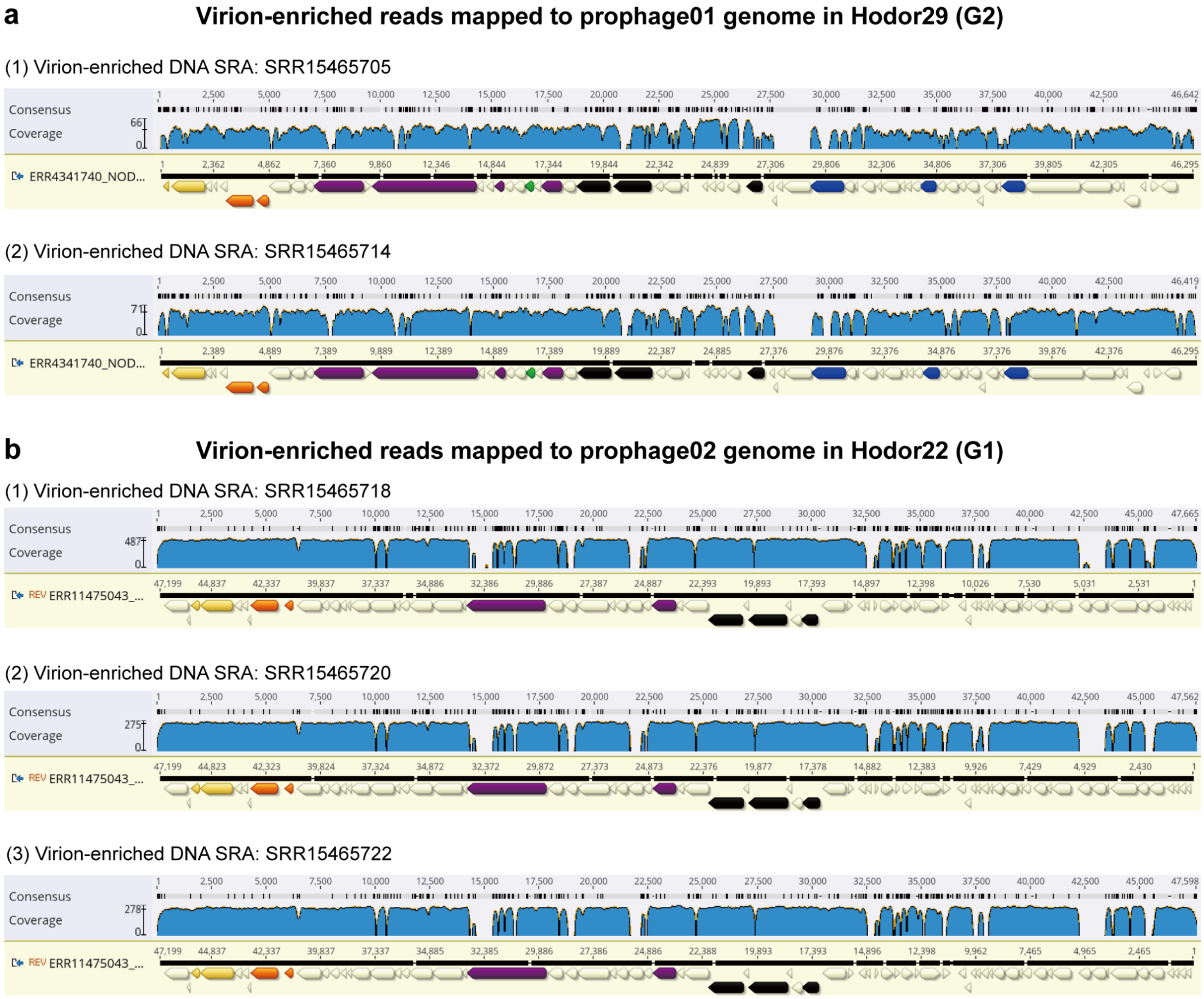


**Supplementary Figure S22 | The mapping profiles of virion-enriched metagenomic reads to prophage genomes in curated Hodor genomes.** (a) Reads from two samples mapped to the prophage01 genome. (b) Reads from three samples mapped to the prophage02 genome. All these reads were from the NCBI project PRJNA755142 [5]. As the reference genomes used for mapping were the prophage genomes in the curated Hodor genomes, instead of those reconstructed from the virion-enriched samples, the profiles have some regions with no reads mapped. As prophage03-related sequences were only detected in 19 Logan-assembled samples, and none of them were virion-enriched (see Supplementary Table 14 for details), we thus do not show them here. The mapping was performed using Bowtie2, and visualized in Geneious Prime Build 2025-03-24 [2].
